# Gut Microbes and the Liver Circadian Clock Partition Glucose and Lipid Metabolism

**DOI:** 10.1101/2022.05.24.491361

**Authors:** Katya Frazier, Sumeed Manzoor, Katherine Carroll, Orlando DeLeon, Sawako Miyoshi, Jun Miyoshi, Marissa St George, Alan Tan, Mariko Izumo, Joseph S. Takahashi, Mrinalini C. Rao, Vanessa A. Leone, Eugene B. Chang

**Affiliations:** Department of Medicine, The University of Chicago, Chicago, IL 60637; Committee on Molecular Metabolism & Nutrition, The University of Chicago, Chicago, IL 60637; Department of General Medicine, Kyorin University School of Medicine, Tokyo, Japan 1818611; Department of Gastroenterology and Hepatology, Kyorin University School of Medicine, Tokyo, Japan 1818611; Department of Neuroscience, University of Texas Southwestern Medical Center, Dallas, TX 75390; The Howard Hughes Medical Institute, University of Texas Southwestern Medical Center, Dallas, TX 75390; Department of Animal & Dairy Sciences, University of Wisconsin-Madison, Madison, WI, 53706

**Keywords:** *Bmal1*, liver, gut microbiota, oscillations, gluconeogenesis, lipid oxidation, tolerance test, RNA- seq, fuel utilization

## Abstract

Circadian rhythms govern glucose homeostasis, and their dysregulation leads to complex metabolic diseases. Gut microbes also exhibit diurnal rhythms that influence host circadian networks and metabolic processes, yet underlying mechanisms remain elusive. Here, we show hierarchical, bi-directional communication between the liver circadian clock, gut microbes, and glucose homeostasis in mice. The liver clock, but not the forebrain clock, requires gut microbes to drive glucose clearance and gluconeogenesis. Liver clock dysfunctionality expands proportions and abundances of oscillating microbial features by two-fold relative to controls. The liver clock is the primary driver of differential and rhythmic hepatic expression of glucose and fatty acid metabolic pathways. Absent the liver clock, gut microbes provide secondary cues that dampen these rhythms, resulting in reduced utilization of lipids as fuel relative to carbohydrates. Together, the liver clock transduces signals from gut microbes necessary to regulate glucose and lipid metabolism and meet energy demands over 24 hours.

**Highlights:**

The liver circadian clock is autonomous from the central clock in metabolic regulation
Liver clock and gut microbes interact to direct hepatic glucose and lipid metabolism
Reciprocating host-microbe interactions drive rhythmic hepatic transcription
Perturbed liver *Bmal1* results in chaotic downstream oscillators and metabolism

## Introduction

Circadian rhythms are essential to nearly all life forms, coordinating behavior and compartmentalizing key physiological processes that govern energy balance efficiency over a 24-hour period (Li et al., 2012). Self-sustaining and cell-autonomous, circadian rhythms serve to synchronize metabolism across the active and inactive phases of life (Sancar and Brunner, 2014), including the transition between fed and fasted states which is a key component of glucose regulation (Berg et al., 2002). Circadian disruption has been linked to metabolic syndrome (Maury et al., 2014), and glucose dysregulation is a hallmark of the disease (Chung et al., 2015; König et al., 2012). Gluconeogenesis (GNG), the endogenous production of glucose, is critical during periods of prolonged fasting (Sharabi et al., 2015). Along with the significant influence of endogenous hormones (Kraus-Friedmann, 1984), both circadian rhythms and gut microbial cues drive GNG, and disruption of either leads to aberrant hepatic GNG (Krisko et al., 2020; Zhang et al., 2010). Few mechanistic insights explain how these consequences arise, thus understanding how these systems function in the context of metabolic homeostasis is of utmost importance.

The circadian clock molecular components are expressed in nearly all cells, forming an elegant feedback loop that organizes gene expression (Buhr and Takahashi, 2013; Takahashi, 2017). *Bmal1* and *Clock* serve as the positive arm that activates gene transcription, including *Cryptochrome (Cry) 1-*2 and *Period (Per) 1-3* which serve as the negative arm to repress *Bmal1* and *Clock*. This results in subsequent reduced expression of *Cry1-*2 and *Per1-3*, allowing the positive arm to resume expression (Ko and Takahashi, 2006). Following genetic disruption of clock genes, metabolic networks become imbalanced, leading to disorders such as glucose intolerance and insulin resistance (Evans and Davidson, 2013). While the central brain clock serves as the master regulator, peripheral tissue clocks exhibit unique oscillations and their specific disruption results in differential metabolic outcomes (Christou et al., 2019; Harfmann et al., 2016; Sadacca et al., 2011; Shimba et al., 2011; Yu et al., 2021). For instance, mice deficient in hepatic *Bmal1* uniquely display increased fasted glucose clearance, altered expression of hepatic GNG genes, and reduced functionality of mitochondria (Lamia et al., 2008; Jacobi et al., 2015).

The gut microbiome is a key regulator of global host metabolism. Microbes vitally contribute to digestion and play a significant role in programming host energy balance (Bäckhed et al., 2007; Martinez-Guryn et al., 2018; Turnbaugh et al., 2006). Their importance in metabolic homeostasis is established through the use of germ-free (GF) mice raised in complete absence of microbes (Bäckhed et al., 2004). Environmental changes, such as diet, can rapidly affect microbial composition and functions that feed back onto the host (Devkota et al., 2012; Huang et al., 2013; Martinez-Guryn et al., 2018). Disturbances in sleep via jet-lag also alter gut microbiome composition in both mouse and human models (Thaiss et al., 2014; Voigt et al., 2014; Benedict et al., 2016). Indeed, gut microbes are essential for normal energy balance and influence liver metabolic function, including hepatic GNG and glucose tolerance (Krisko et al., 2020; Krajmalnik-Brown et al., 2012; Wichmann et al., 2013).

Gut microbes provide critical inputs that drive host circadian rhythms and metabolism, and exposure to high-fat diet disturbs these rhythms and functionality in mice (Leone et al., 2015; Murakami et al., 2016). High-fat diet also disrupts diurnal oscillations of microbial abundance and metabolite levels, while timed feeding can somewhat recover this effect (Eggink et al., 2017; Leone et al., 2015; Zarrinpar et al., 2014). Mice with global genetic mutations in circadian clock genes exhibit significantly altered microbial community profiles and loss of oscillations in specific taxa (Brooks et al., 2021; Liang et al., 2015; Thaiss et al., 2014). Conversely, in absence of gut microbial cues, GF wild-type mice exhibit dampened rhythmicity in the expression of core circadian clock genes in both brain and liver, as well altered transcriptomic patterns in liver, small intestine, and white adipose tissue (Leone et al., 2015; Weger et al., 2019). While the link between circadian rhythms, gut microbiota, and host metabolism is established, few mechanisms have been proposed to explain how these phenomena occur, how the circadian clock in specific metabolic organs such as the liver are involved, and how this impacts global metabolic regulation. We hypothesized that hepatic GNG is driven by bidirectional interactions between the hepatic circadian clock and gut microbiota.

Here, using the liver-specific *Bmal1* knock-out mouse raised in both conventional and GF conditions, we demonstrate a hierarchy of signals between the gut microbiome and liver clock that coordinates hepatic GNG and fatty acid β-oxidation (FA β-Ox) that are essential to maintain diurnal rhythms in glucose homeostasis. We identify that regardless of microbial status, the liver clock is a primary driver of hepatic transcription, particularly of genes involved in the organization and function of key metabolic pathways. Secondarily, the liver clock transduces timed signals from the gut microbiome to promote coordinated differential, oscillatory, and correlated transcription patterns in glucose and lipid metabolic pathways. When either of these two components (host clock or microbes) are absent or dysfunctional, we observe a significant gain in oscillating hepatic transcripts that are disorganized. The absence of a functional clock also results in expansion of oscillations in specific fecal microbial abundances. Together, we reveal interactions between the liver circadian clock and gut microbes that aid in glucose and lipid homeostasis that govern whole-body metabolism and fuel utilization.

## Results

### Gut microbes are essential for liver circadian clock regulation of insulin-independent glucose clearance

To assess the effects of a dysfunctional liver clock and gut microbes on mammalian host metabolism, we utilized male and female mice with liver-specific deletion of *Bmal1* gene expression via *Albumin*-*cre*: *Bmal1^f/f^ x Albumin*-*cre* (LKO) versus control, *Bmal1^f/f^*(WT) (Postic et al., 1999; Storch et al., 2007). We confirmed *Bmal1* deletion in the liver, while expression in the brain remained intact (**Figure S1A**). To assess the role of microbiota, we maintained both WT and LKO mice in conventional, specific-pathogen free (SPF) or germ-free (GF) conditions.

First, we observed that SPF LKO male mice exhibited significantly increased body weight compared to SPF WT, while oppositely GF LKO mice exhibited a slight, non-significant trend of decreased body weight relative to GF WT (**Figure 1A**, upper panel). This difference in body weight could not be explained by changes in gross liver or fat pad tissue weight (**Figure S1B**). Weekly caloric consumption revealed similar patterns; SPF LKO mice ate slightly but significantly more than WT, while GF LKO mice ate slightly but significantly less than their WT counterparts (**Figure 1A**, lower panel). Interestingly, we did not observe any differences between genotype in female mice for body weight or food intake (data not shown). We then measured resting blood glucose levels every 4 hours over a 12:12LD cycle and found that overall levels were not drastically different between groups in male mice (**Figure 1B**). However, analysis by CircWave revealed only WT mice, regardless of microbial status, exhibited significant oscillation of blood glucose levels, indicating that diurnal circulating blood glucose is driven by the liver clock and not gut microbes. Conversely, we observed very little difference in diurnal blood glucose values in female mice (data not shown).

**Figure 1.**
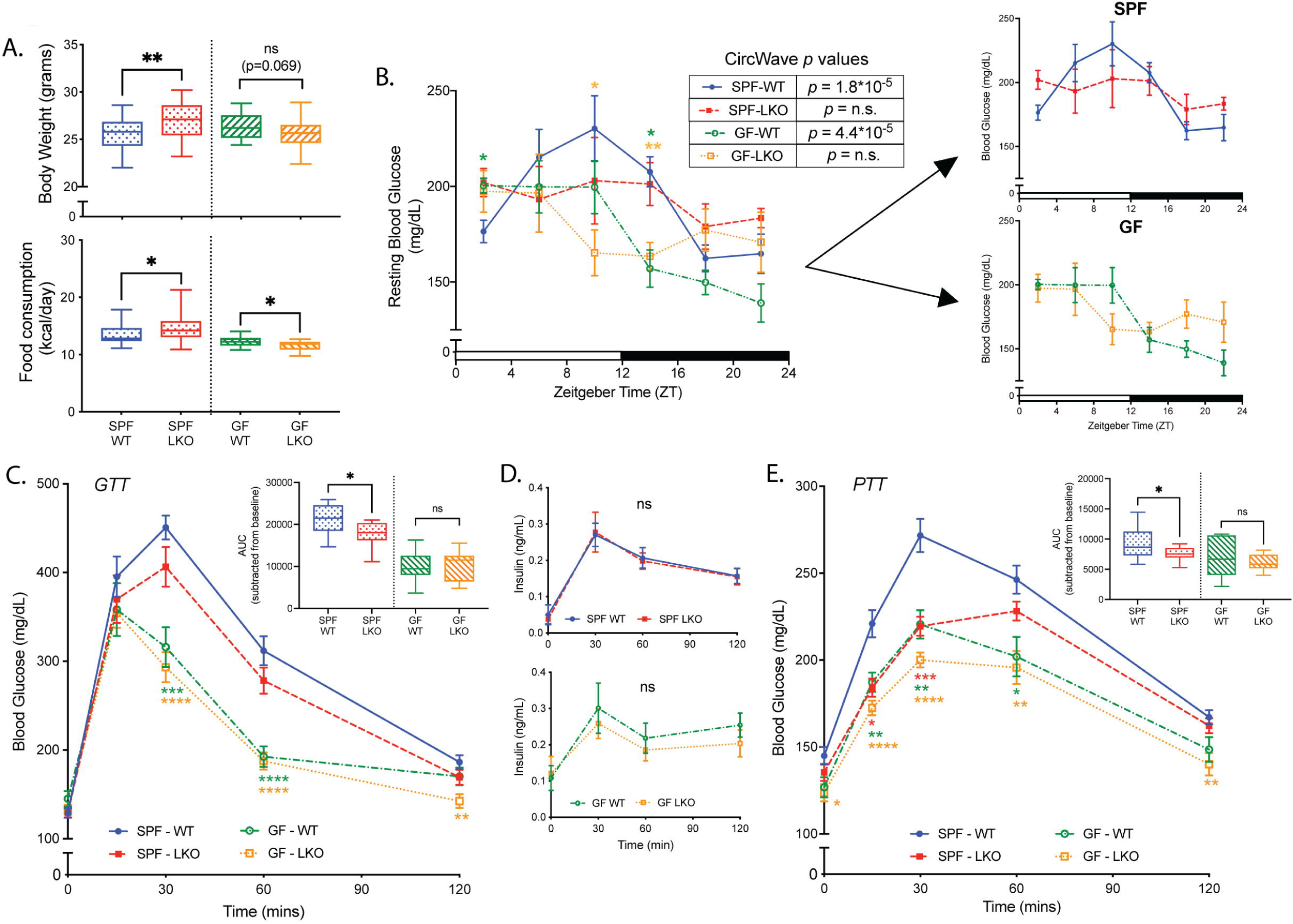
Gut microbes are essential for liver circadian clock-mediated glucose metabolism. **(A)** Body weight (upper panel) and weekly food intake (lower panel) from SPF and GF, WT and LKO male mice (*n*=18-32/group). **(B)** Resting blood glucose levels measured via tail snip of SPF and GF, WT and LKO male mice every 4 hrs over 24 hrs (*n*=4-6/group/timepoint, SPF and GF groups also shown separately). CircWave statistics indicate significantly oscillating (p<0.05) or not oscillating (p>0.05) values. **(C)** Oral Glucose Tolerance Test of SPF and GF, WT and LKO male mice (GTT, *n*=10-13/group). **(D)** Circulating insulin levels during GTT (*n*=10-13/group). **(E)** Intraperitoneal Pyruvate Tolerance Test (PTT, *n*=10-15/group) of SPF and GF, WT and LKO male mice. Data points represent mean±SEM, box plots represent median±min/max. ****p<0.0001, ***p<0.001, **p<0.01, *p<0.05, ns=not significant; colored stars represent significance relative to SPF WT. Inset graphs represent area under the curve (AUC) normalized to baseline glucose.

Given previous work by Lamia *et al*. (2008), we performed an oral glucose tolerance test (GTT) and observed that male GF mice exhibit more rapid glucose clearance than SPF by area under the curve (AUC) (**Figure 1C**). We also observed a genotype-driven effect in SPF conditions where LKO mice cleared glucose significantly faster than WT. Circulating insulin levels during GTT were not significantly different between WT and LKO mice in either SPF or GF conditions, suggesting that insulin secretion is not impacted by liver clock functionality (**Figure 1D**). GTT of female mice revealed no significant differences in AUC between any group (**Figure S1C**), suggesting that liver-clock-mediated glucose clearance is in part driven by sex. We then interrogated insulin sensitivity by intraperitoneal insulin tolerance test (ITT) on SPF and GF, WT and LKO male mice and revealed no difference in insulin sensitivity between genotypes in either microbial condition (**Figure S1D**), suggesting that male *Bmal1*-driven glucose clearance is insulin-independent.

These data suggest the liver circadian clock imparts an insulin-independent effect on glucose clearance in male mice that is dependent on the presence of gut microbes.

### Circadian clock and microbiome-driven GNG is liver-specific and requires *in vivo* signals

To interrogate the role of the liver clock on GNG, we performed an intraperitoneal pyruvate tolerance test (PTT). First, we revealed GF mice exhibit significantly lower GNG rates than SPF counterparts regardless of genotype (**Figure 1E**). Second, we observed LKO mice exhibit reduced GNG in SPF conditions, while no differences between genotypes in GF conditions were evident. Additionally, we performed PTT of female SPF WT and LKO mice and observed no difference in GNG (**Figure S1E**). Following our results that liver-clock-mediated glucose clearance and GNG is male-specific, we proceeded with only male mice for the duration of the study.

To determine whether reduced GNG is liver-clock-specific, we performed PTT on mice lacking a functional core brain clock by *CamkIIa*-*cre* forebrain-specific *Bmal1* knockout, while peripheral clocks remain intact (Casanova et al., 2001; Izumo et al., 2014) (**Figure S1A**). We found no difference in GNG rates between WT and forebrain-*Bmal1*-KO mice (**Figure S1F**), indicating that clock-mediated changes in GNG are specific to hepatic *Bmal1* and liver clock. This result corroborates previous evidence that the liver clock, specifically hepatic *Bmal1*, is somewhat autonomous from the core clock located in the brain (Koronowski et al., 2019).

Since both GNG and glycogenolysis are co-stimulated during periods of prolonged fasting, we measured glycogen levels in liver samples collected from each group every 4 hrs over 24 hrs. We observed no difference between SPF groups at any timepoint, and glycogen content was only elevated in GF WT relative to LKO mice at ZT22 (4am) (**Figure S1G**). In all groups, liver glycogen levels exhibited significant and remarkably similar oscillations over a 24hr period, as evident by CircWave statistics. These data indicate that hepatic glycogenolysis is not a major contributor to either the microbe- or clock-mediated effect on the observed glucose homeostasis phenotype, further supporting that GNG is the major contributor.

Finally, we examined glucose production in primary hepatocytes isolated from SPF and GF WT vs LKO *ex vivo* following stimulation with GNG-inducing substrates in glucose-free media. We revealed no difference in glucose production in media between groups following GNG stimulation with a cell permeable cAMP analog (cPT-cAMP) (**Figure S1H**). This suggests the cellular machinery to perform GNG is not affected by liver clock functionality or prior exposure to gut microbes, and additional *in vivo* signals are necessary for liver clock- and microbe-mediated GNG.

Overall, we reveal a microbiota-dependent effect of liver circadian clock-mediated GNG that requires real-time *in vivo* signals.

### Gut microbes are necessary, but not sufficient, for liver clock-mediated GNG

To investigate the role of gut microbes, we performed PTT in adult SPF WT and LKO mice both before (Pre-Abx) and after (Post-Abx) acute elimination of gut microbes via daily antibiotic treatment for 2 weeks. We also confirmed significant bacterial reduction by 16S gene copy number in Post-Abx stool (**Figure S2A**). While PTT of Pre-Abx mice mimicked our SPF results in **Figure 1E**, we observed no differences in GNG between genotypes Post-Abx, closely resembling outcomes in GF mice (**Figure 2A**). This supports the hypothesis that gut microbial cues are essential for *in vivo* coordination of GNG mediated by the liver clock.

**Figure 2.**
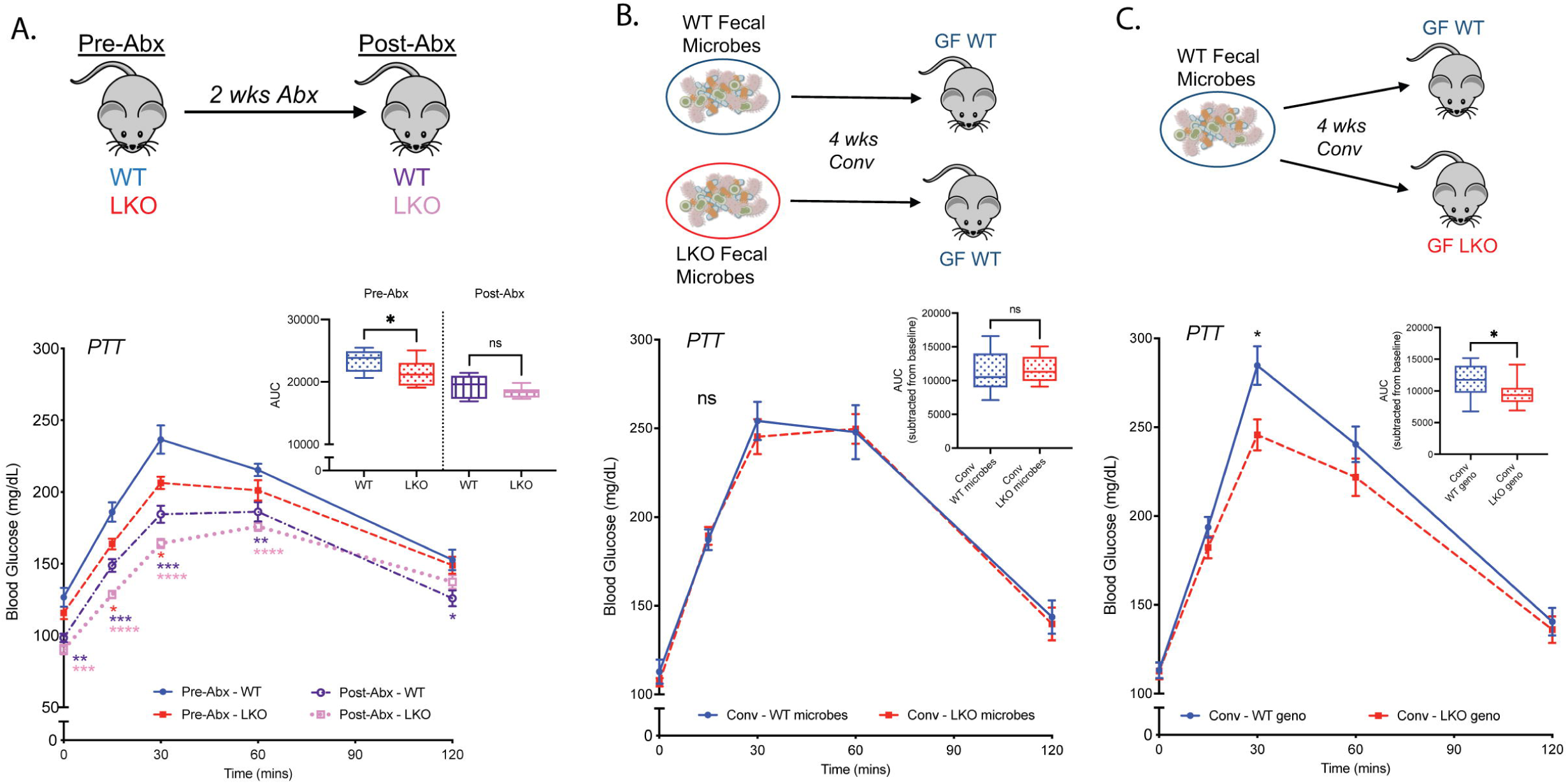
Modulation of gut microbes can both eliminate and restore liver-clock mediated GNG. **(A)** PTT in WT and LKO male mice before (Pre-Abx) and after (Post-Abx) daily antibiotic treatment for 2 weeks (n=12-13/group); inset graph represents AUC. **(B)** PTT in GF WT male mice conventionalized with fecal microbes from SPF WT or LKO male mice (n=11-12/group). **(C)** PTT in GF WT and LKO male mice conventionalized with fecal microbes from SPF WT male mice (n=15-16/group); inset graphs represent AUC normalized to baseline glucose. Data points represent mean±SEM, box plots represent median±min/max. ****p<0.0001, ***p<0.001, **p<0.01, *p<0.05, ns=not significant; colored stars represent significance relative to Pre-Abx WT.

Next, we conventionalized adult GF C57Bl/6J WT mice via fecal microbiota transplant from either WT or LKO SPF donor mice and performed PTT. Here, we did not observe significant differences between recipient groups (**Figure 2B**). This suggests gut microbes selected by the presence or absence of a functional liver clock alone were insufficient to transfer the GNG phenotype, implying that genotype is the primary driver of GNG. We then performed a different microbiota transplant in which we conventionalized GF WT or LKO recipient mice with identical WT donor fecal microbes (**Figure 2C**). We observed significantly reduced GNG in LKO mice relative to WT recipients, indicating that restoration of the LKO SPF phenotype requires both gut microbes and genetic absence of a liver clock.

Given that gut microbes directly modulate GNG via the liver clock, we sought to determine if loss of *Bmal1* impacts microbiota community membership. Co-housing mice can normalize microbiota differences and mask the impact of genotype, largely due to the coprophagic tendencies of mice, so we compared SPF WT and LKO mice in a mixed housing scenario (WT + LKO) with mice separated by genotype at weaning (WT + WT vs. LKO + LKO) (**Figure S2B**). Stool was collected at 12 weeks of age for Illumina Mi-Seq 16S rRNA gene sequencing. Regardless of housing, we detected no significant differences in overall community membership via Beta-diversity metrics (data not shown) or by relative abundance of amplicon sequence variants (ASVs) belonging to dominant phyla (**Figure S2C**) or less abundant phyla (**Figure S2D**).

Taken together, liver clock functionality does not impact overall gut microbiota community assemblage but serves as the primary driver of GNG, while microbes are key secondary drivers. This suggests a concurrent interaction between gut microbes and the liver clock that requires real-time signals from gut microbes to mediate GNG.

### The liver clock drives oscillations of specific microbiota community members

Global murine deletion of *Bmal1* has previously been shown to alter diurnal oscillations of fecal microbiota (Liang et al., 2015). Thus, we inquired whether liver clock functionality impacted diurnal oscillations of fecal microbiota collected from WT and LKO male mice every 6 hrs over 48-hrs (**Figure S3A**). We found that bacterial load was similar between WT and LKO across all timepoints (**Figure S3B**). We next employed empirical Jonckheere-Terpstra-Kendall CYCLE (eJTK) to identify significantly oscillating ASVs and discovered that LKO mice exhibit nearly twice the number of oscillating ASVs compared to WT (**Figure 3A, S3C**). The influx of unique oscillating microbes in LKO was mostly attributed to ASVs identified to class Clostridiales. Conversely, we observed little difference in the number of oscillating Bacteroidales ASVs, suggesting LKO may specifically impact diurnal oscillations of Clostridiales microbes.

**Figure 3.**
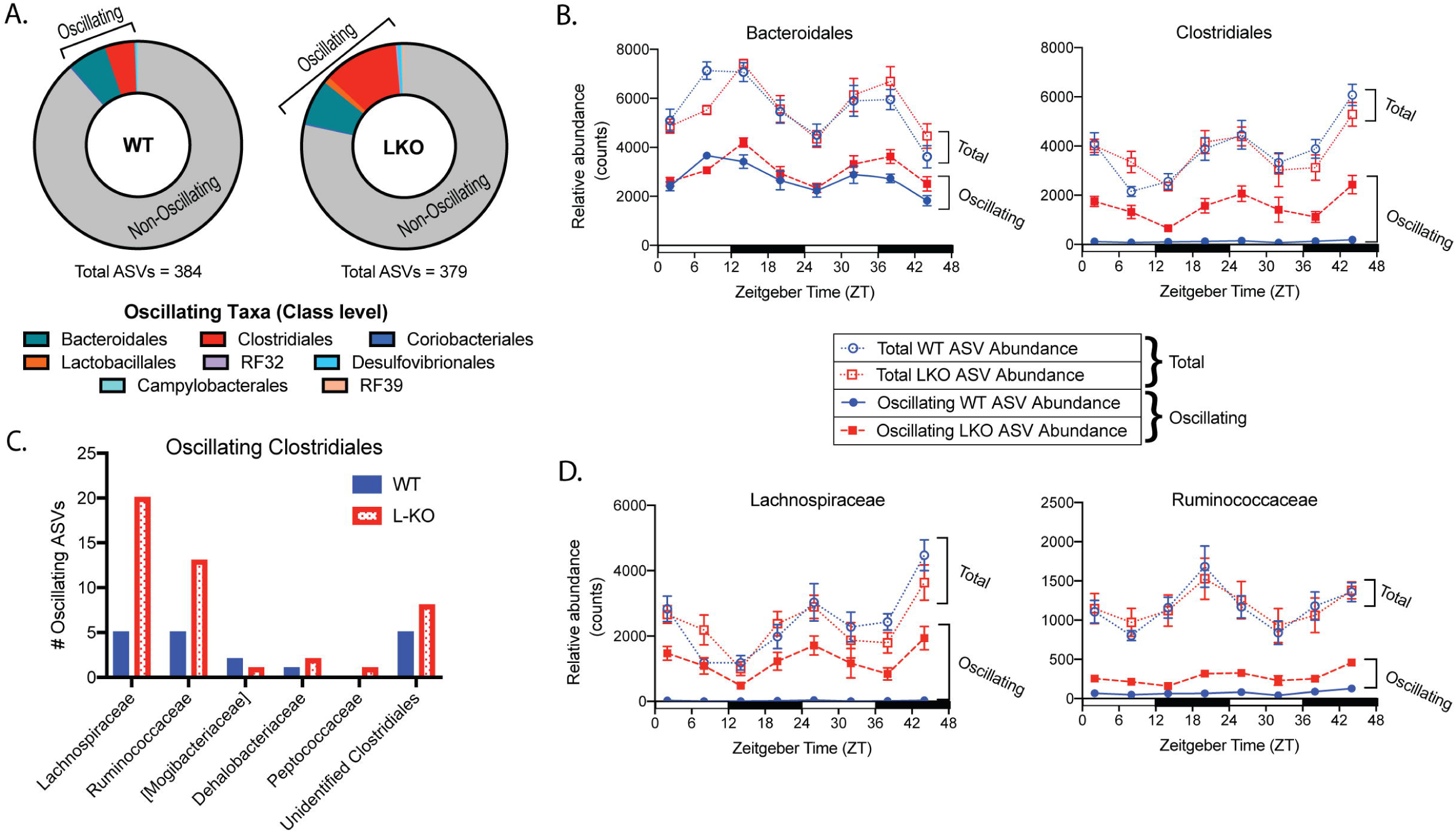
Liver circadian clock drives unique patterns of oscillations in microbial abundance. 16S rRNA gene sequencing of stool sampled from SPF WT and LKO male mice every 6 hrs over 48-hrs via repeat-collection (*n*=7-8/group). **(A)** Proportion of non-oscillating (grey area) vs significantly oscillating (colored areas) Amplicon Sequence Variants (ASVs) identified via eJTK; oscillating ASVs divided by taxonomic class. **(B)** Relative abundance counts of total vs oscillating ASVs within classes Bacteroidales and Clostridiales. **(C)** Number of oscillating Clostridiales ASVs at the family level in WT and LKO. **(D)** Relative abundance counts of total vs oscillating ASVs within families Lachnospiraceae and Ruminococcaceae. Data represent mean±SEM.

Aside from proportion, the relative abundance of oscillating ASVs was also significantly greater in LKO relative to WT (**Figure S3D**). Upon plotting abundance of total vs oscillating Bacteroidales and Clostridiales ASVs, we identified that overall abundance was not different between WT and LKO, but only Clostridiales exhibited increased abundance of oscillators in LKO (**Figure 3B**). We then examined oscillating Clostridiales at the family level and identified that Lachnospiraceae and Ruminococcaceae accounted for the gain of uniquely oscillating ASVs in LKO (**Figure 3C**). While the total abundance of either family was not different between genotypes, the abundance of oscillating ASVs significantly increased within LKO (**Figure 3D**). Interestingly, among the ASVs annotated to species, *Lactobacillus murinus* followed the same pattern of gained oscillation in LKO while total abundance remained constant (**Figure S3E**).

Despite no changes in overall microbiota abundance profiles, the loss of a functional liver clock drives a unique signature of increased diurnal oscillations in specific stool microbiota community members, particularly those belonging to the class Clostridiales.

### The liver clock is the main driver of hepatic transcriptomes, secondarily influenced by gut microbes

Given that liver clock and gut microbes regulate GNG, we sought to determine if other metabolic pathways were impacted. We performed RNA-sequencing of liver samples collected every 4 hours over 24-hours from SPF and GF, WT and LKO male mice, and the data were analyzed via three approaches (**Figure 4A**). Principle Component Analysis of pooled samples revealed distinct separation by genotype along PC1 and microbial status along PC2 (49% and 12% of variance, respectively) (**Figure 4B**). Similar patterns were observed when samples were divided by timepoint, demonstrating consistency across a 24-hour period (**Figure S4A**). This suggests a hierarchy in which liver clock functionality (genotype) is the primary driver of the hepatic transcriptome, while a gut microbiome serves as a secondary driver.

**Figure 4.**
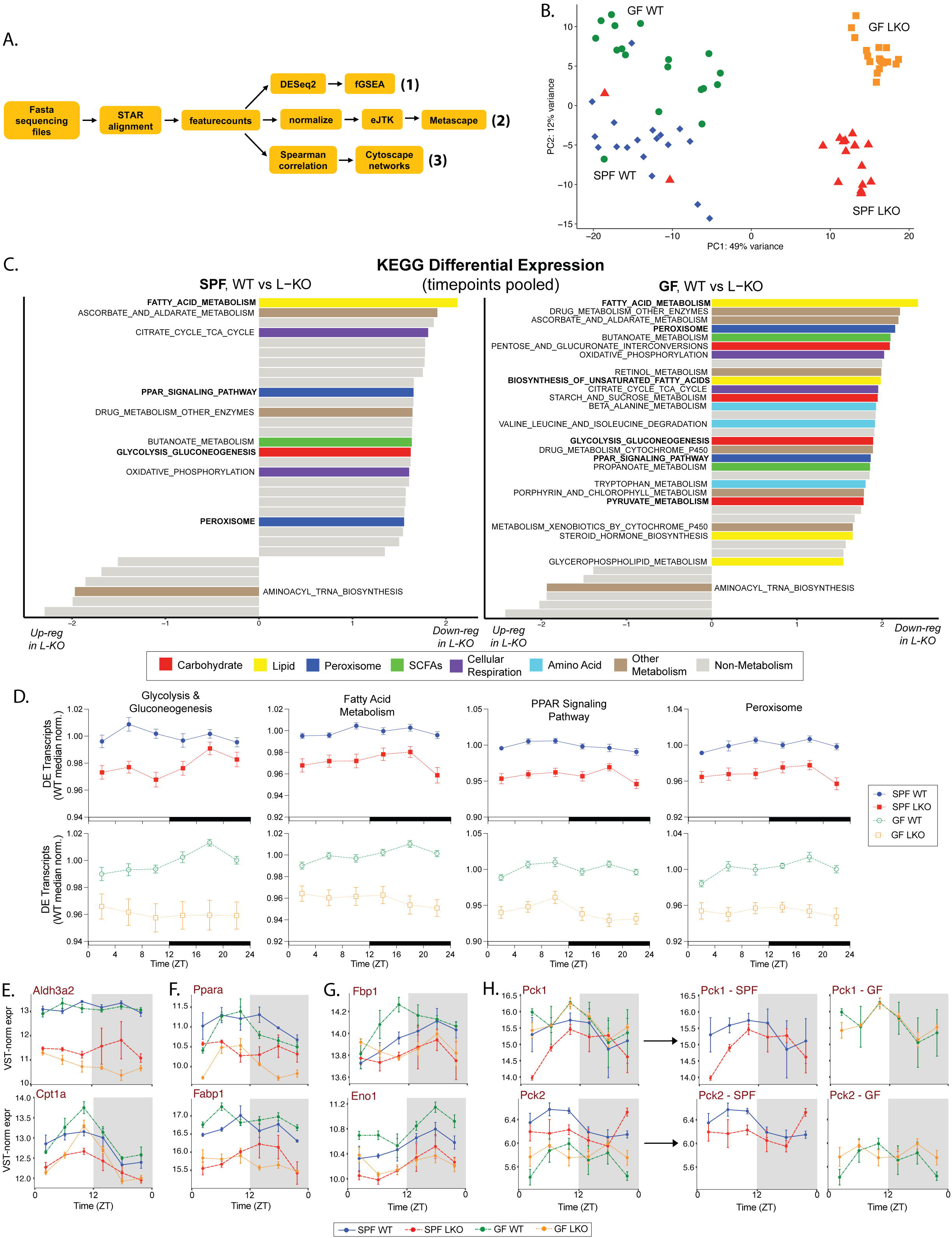
Metabolic pathway gene expression is downregulated in absence of a liver clock across time and regardless of microbial status. Transcriptome analysis of liver samples collected every 4 hrs over 24-hrs (ZT2, 6, 10, 14, 18, 22) from SPF and GF, WT and LKO male mice (n=3/timepoint/group). **(A)** Data analysis workflow, demonstrating three arms of analysis: 1. differential expression (Fig 4C-H, S4), 2. oscillation (Fig 5, S5A), and 3. network co- occurrence (Fig 6, S5B-E). **(B)** Principal Component Analysis of transcriptome profiles, all samples pooled. **(C)** Differentially regulated KEGG pathways between WT and LKO, within SPF and GF, timepoints pooled. Metabolic pathways colored according to legend, non-metabolic pathways colored gray. Bars to the right of plot midline represent pathways downregulated in LKO compared to WT; bars to the left represent pathways upregulated in LKO compared to WT. Bold pathway titles are addressed intext. **(D)** WT-median-normalized expression of differentially expressed (DE) genes within identified KEGG pathways. **(E-H)** VST-normalized expression of leading-edge genes in FA metabolism **(E)**, PPAR signaling **(F)**, and GNG **(G,H)** differentially expressed between WT and LKO in SPF and GF, with the exception of *Pck1* & *Pck2* (H) which are only differentially expressed between SPF genotypes (SPF and GF groups also shown separately). Data represent mean±SEM.

First, we performed differential gene expression analysis via DESeq2 and fast Gene Set Enrichment Analysis (fGSEA) to identify molecular pathways impacted by the liver clock and/or gut microbes (**Figure 4A**, Approach 1). Analysis using Kyoto Encyclopedia of Genes and Genomes (KEGG) revealed nearly all differentially expressed metabolic pathways were downregulated in LKO compared to WT, regardless of microbial status (**Figure 4C**). However, GF conditions exhibited more downregulated metabolic pathways in LKO than SPF conditions. This may suggest that gut microbes stabilize pathway transcriptional regulation; that is, certain *Bmal1*-driven effects only emerge in absence of microbes. Differential analysis within collection timepoint revealed nearly identical patterns of overall downregulation of metabolic pathways (**Figure S4B**). Importantly, we observed downregulation of “Glycolysis_Gluconeogenesis” in SPF LKO compared to WT (**Figure 4C**, left panel), confirming our evidence that GNG is impaired in SPF LKO (**Figure 1E**). We also observed downregulation of GNG pathways, including “Pyruvate_Metabolism”, in GF LKO compared to WT (**Figure 4C**, right panel), while our physiological evidence reveals no difference in overall GNG output between GF WT and LKO (**Figure 1E**). This suggests that while differences in transcription between genotypes are similar in GF and SPF, additional signals contribute to reduced GNG in GF conditions regardless of genotype.

Apart from GNG, we observed LKO downregulation of several metabolic pathways involved in FA and lipid metabolism, including “Fatty_Acid_Metabolism”, “PPAR_Signaling_Pathway”, “Peroxisome”, and “Biosynthesis_of_Unsaturated_Fatty_Acids” (**Figure 4C**). This corroborates previous findings that lipid metabolic gene expression is downregulated in SPF LKO mice compared to WT via RNA methylation (Zhong et al., 2018). The regulation of GNG and FA metabolism are intricately tied; under fasting conditions, FA β-Ox is activated to provide acetyl-CoA to generate ATP, which sustains the conversion of pyruvate and other GNG substrates into glucose (Jones, 2016). Thus, the regulation of these two processes is closely linked and changes in one directly impact the other (Randle, 1998).

Given this differential regulation, we plotted median-normalized expression for all leading-edge transcripts within relevant pathways, confirming significant down-regulation in LKO compared to WT in both SPF and GF (**Figure 4D, S4C**). We next examined expression of specific transcripts within these pathways, in particular key transcripts involved in FA metabolism (**Figure 4E**; *Aldh3a2:* Aldehyde dehydrogenase 3 family member A2*; Cpt1a:* Carnitine palmitoyltransferase 1A), PPAR signaling (**Figure 4F**; *Ppara:* Peroxisome proliferator activated receptor alpha*; Fabp1:* Fatty acid binding protein 1), and GNG (**Figure 4G,H**; *Fbp1:* Fructose-bisphosphate 1*; Eno1:* Enolase 1*; Pck1/2:* Phosphoenolpyruvate carboxylase 1/2). Expression was significantly reduced in many of these transcripts in LKO compared to WT regardless of microbiota status (**Figure 4E-G**). However, expression of the key rate-limiting GNG enzymes *Pck1* and *Pck2* exhibited more nuanced patterns (**Figure 4H**). *Pck1* revealed significantly reduced expression in SPF LKO compared to WT, while both GF groups exhibited expression levels similar to SPF WT. Conversely, *Pck2* was significantly reduced in SPF LKO compared to WT, and both GF groups were reduced compared to SPF, mirroring our PTT results (**Figure 1E**). Thus, *Pck2* may be key for liver-clock- and microbially-mediated effects on GNG, while *Pck1* is only different under SPF conditions.

In summary, the liver clock is the primary driver of transcriptome differential regulation, particularly for key metabolic pathways, such as GNG, that are downregulated following loss of liver clock. However, gut microbes provide additional cues to modulate expression, including LKO-driven expression of the rate-limiting GNG enzyme *Pck2*.

### Liver clock and gut microbes drive unique hepatic transcriptome oscillations

Following previous studies (Thaiss et al., 2016), we identified all significantly oscillating transcripts within each group via eJTK (**Figure 4A**, Approach 2). SPF LKO exhibited more oscillating transcripts than WT (1503 vs 1104), and GF LKO exhibited a similar increase relative to GF WT (3580 vs 2544) (**Figure 5A**). Only 155 transcripts were oscillating in all groups, while many were uniquely oscillating in a single group. This emergence of unique oscillators in the absence of a functional liver clock mirrored the pattern observed in ASV relative abundance in repeat-collected stool (**Figure 3A**). These data reveal that loss of key drivers (i.e., the liver clock or gut microbes) results in emergence of unique oscillatory elements both in the hepatic transcriptome and gut microbiota community, which may contribute to a loss in metabolic homeostasis *in vivo*.

**Figure 5.**
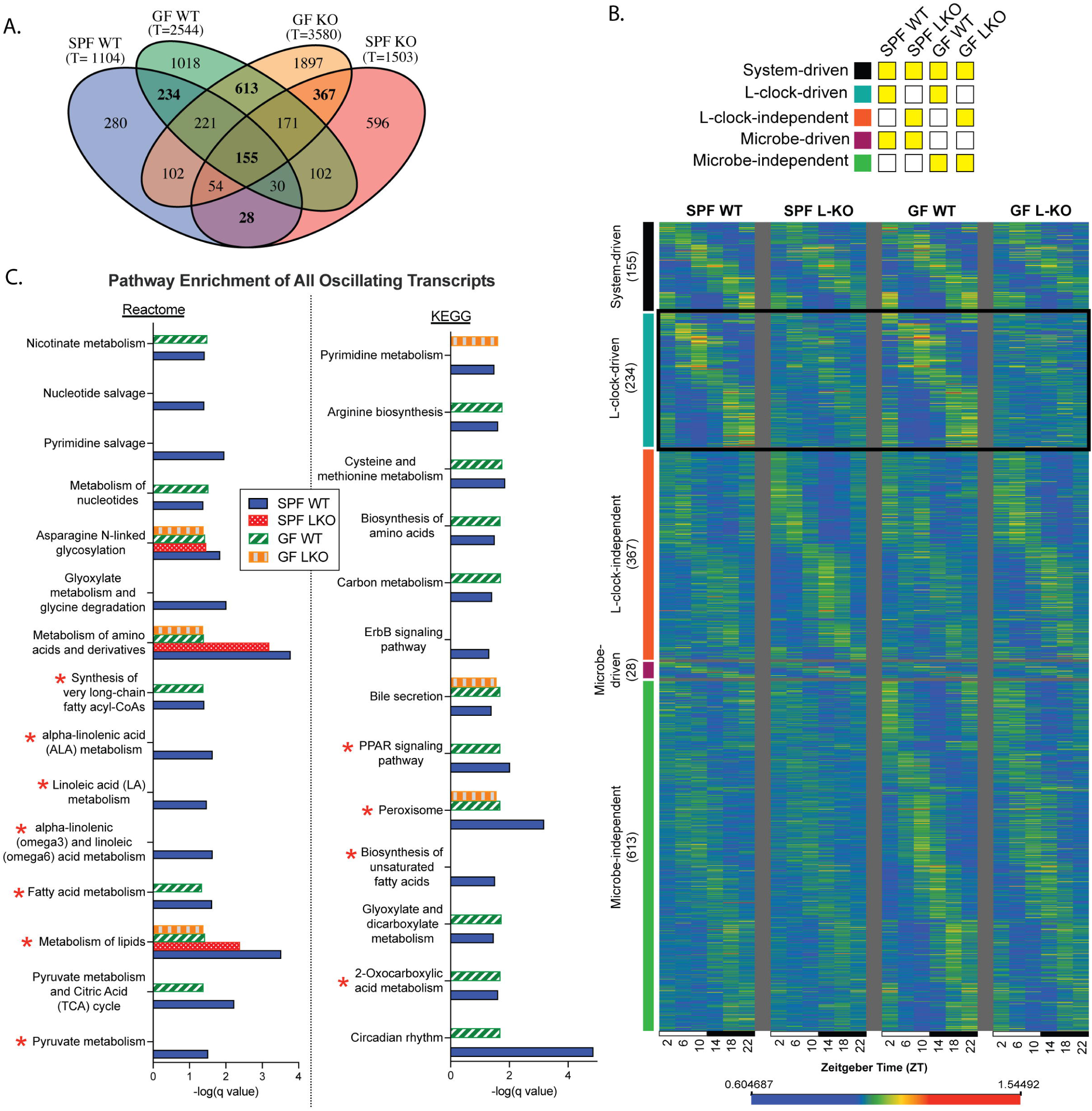
The liver clock and gut microbes drive unique hepatic transcriptome oscillations, affecting functional pathway enrichment. Diurnal transcriptome analysis of liver samples collected every 4 hrs over 24-hrs from animals maintained in 12:12 LD (ZT2, 6, 10, 14, 18, 22) from SPF and GF, WT and LKO male mice (n=3/timepoint/group). **(A)** Venn diagram of significantly oscillating transcripts across each group; total number of oscillating transcripts are under each group title; oscillating transcripts identified via eJTK (GammaBH<0.05); bold numbers are visualized in Fig 5B. **(B)** Expression of significantly oscillating transcripts that are system-driven, liver-clock-driven or -independent, and microbe-driven or -independent; expression normalized by median, transcripts ordered by time of max expression and phase; legend indicates which transcripts are depicted in each by yellow highlight. **(C)** Reactome & KEGG pathways significantly enriched by oscillating transcripts within each group; subset of pathways enriched in SPF WT oscillating genes (q<0.05); no bar indicates lack of significance for that group/pathway (q>0.05); pathways marked with red star are addressed intext.

We then compared oscillating gene expression patterns. While SPF WT transcripts exhibit clear diurnal patterns, oscillations were significantly dampened in LKO, regardless of microbial status (**Figure S5A**). Conversely, the oscillation patterns were well-preserved in GF WT, reinforcing that the liver clock is the main driver of core oscillating transcripts. We next partitioned oscillating transcripts to those that are system-driven, liver-clock-driven, liver clock-independent, microbe-driven, and microbe-independent (**Figure 5B**). System-driven transcripts exhibited a conserved oscillation pattern across groups, while the other sets exhibited dampened oscillations. For example, the microbe-independent transcripts demonstrated robust oscillation across GF groups, with clear dampening of oscillation across SPF groups. Interestingly, the only subset of transcripts that exhibited a unique pattern were the liver-clock-driven oscillating transcripts (**Figure 5B**, bolded box). Whereas both WT groups exhibited robust oscillations in liver clock-driven transcripts, LKO exhibited severe dampening. Between LKO groups, SPF LKO displayed a more preserved organization of oscillating transcripts, while the highest level of disorganization was observed in GF LKO relative to WT. This suggests that the liver clock and gut microbes impart combinatorial action on the temporal organization of specific liver-clock-driven diurnal hepatic gene expression.

Overall, the liver clock and gut microbes independently impart a unique impact on oscillating transcripts, and absence of both drivers results in further disorganization of oscillating transcripts.

### Liver-clock and gut microbes exhibit individual and combinatorial influences on hepatic GNG and FA metabolic pathway expression

Next, we applied Metascape (Zhou et al., 2019) to statistically determine which pathways were significantly enriched by each group of oscillating transcripts (q<0.05, summarized in **Table S1**). Examining pathways enriched by SPF WT oscillating liver transcripts relative to other groups, we observed each factor (liver clock and gut microbiota) elicited similar levels, reduced levels, or total absence of enrichment (**Figure 5C**). Both SPF and GF WT exhibited similar enrichment across many pathways, supporting our finding that hepatic transcriptome oscillation patterns are primarily driven by the liver clock. Importantly, we observed enrichment of Reactome “Pyruvate Metabolism” only in SPF WT oscillating transcripts. This loss of oscillating pyruvate metabolism transcripts could contribute to the observed reduction in GNG output detected in SPF LKO and both GF groups (**Figure 1E**). KEGG “2-oxocarboxylic acid metabolism”, which includes pyruvate metabolism, also exhibited loss of enrichment in both LKO groups while WT groups exhibited enrichment (**Figure 5C**).

In addition to glucose metabolic pathways, we also noted differential enrichment of lipid metabolic pathways (**Figure 5C**). Reactome “Metabolism of lipids” was differentially enriched in all groups, where SPF WT exhibited the greatest enrichment, SPF LKO was intermediate, and both GF groups exhibited the lowest enrichment. This signifies a deviation from normal (SPF WT) oscillation patterns in transcripts that are key mediators of lipid metabolism. Importantly, several FA pathways, including “Fatty acid metabolism”, “Biosynthesis of unsaturated fatty acids”, several linoleic acid pathways, “Peroxisome”, and “PPAR signaling pathway” lacked any significant enrichment in SPF LKO oscillating transcripts, and reduced or absent enrichment across both GF groups. The recurrence of altered FA metabolic pathway enrichment, alongside altered pathways associated with GNG expression, supports our previous claim of concurrent and synchronized regulation of these two metabolic pathways that impart key impacts on global glucose regulation.

Together, diurnal oscillations of hepatic gene transcription are significantly altered by absence of a functional liver clock, gut microbes, or both. Although LKO and GF mice exhibit increased, unique oscillating transcripts, the normal enrichment of key GNG and FA metabolic pathways is reduced or lost in LKO and GF, further supporting the impaired efficiency of these metabolic processes *in vivo*.

### Gut microbes impact liver clock-driven network co-occurrence of transcripts

We then examined whether liver clock or gut microbes imposed an impact on transcript correlations over time by co-occurrence network analysis (**Figure 4A**, Approach 3). We calculated Spearman correlation coefficients via pairwise-comparisons of transcript reads over time to identify significant co-occurrences for network visualization (**Figure 6**; network statistics in **Figure S5B**). This allowed for the identification of nodes (correlated transcripts, p<0.001) and their corresponding edges (connections between nodes). SPF LKO exhibited a modest increase in total nodes relative to WT (6116 vs. 5603); however, the number of edges increased from 20,922 (WT) to 36,702 (LKO) (**Figure 6A**), which is visually recognized by the density of the SPF LKO network compared to WT (**Figure 6B**). Conversely, the total number of nodes and edges did not vastly differ between GF WT and LKO. Additionally, the overall density of GF networks, regardless of liver clock status, was greatly reduced in comparison to SPF.

**Figure 6.**
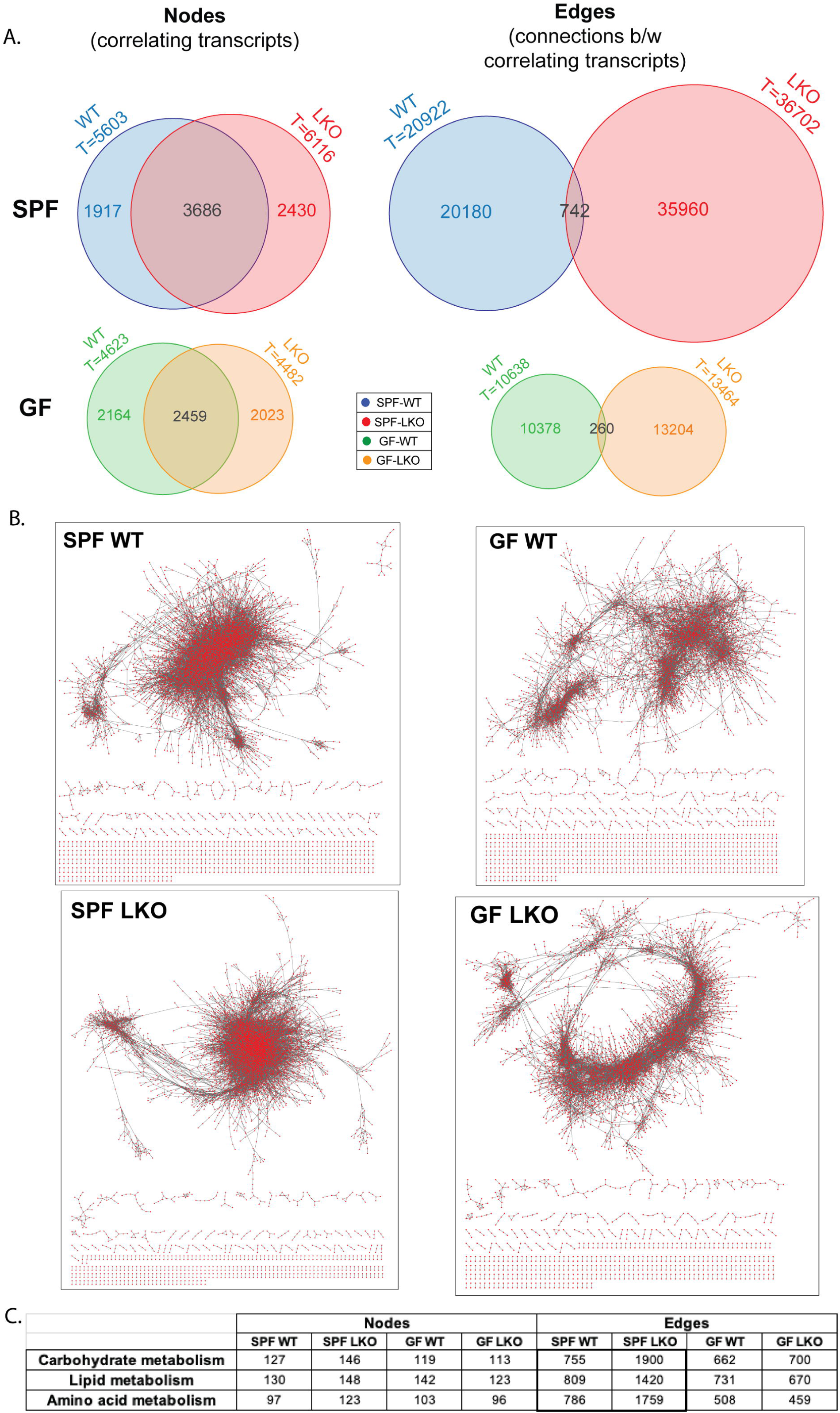
Hepatic transcriptome co-occurrence over time are differentially impacted by the liver clock and gut microbes. Network transcriptome analysis of liver samples collected every 4 hrs over 24-hrs from animals maintained in 12:12 LD (ZT2, 6, 10, 14, 18, 22) from SPF and GF, WT and LKO male mice (n=3/timepoint/group); Network co-occurrence analysis of correlating transcripts over time within each group (p<0.001). **(A)** Venn diagrams summarizing the number of correlating transcripts (nodes) and connections (edges) in each group and crossover within SPF and GF. **(B)** Network visualization; red dots represent nodes, grey lines represent edges. **(C)** Total nodes and edges in each group, annotated to carbohydrate, lipid, and amino acid metabolism by KEGG.

Next, KEGG annotations were applied to significantly correlated nodes and edges. While the number of nodes annotated to Carbohydrate (KO09101), Lipid (KO09103), and Amino acid (KO09105) metabolic pathways were not vastly different between groups, SPF LKO exhibited a two-fold increase in edges annotated to these pathways compared to SPF WT (**Figure 6C** bolded box, **S5C-E**), while no difference was observed between GF groups. Despite a modest influence of liver clock or gut microbes on the total number of connected transcripts, loss of functional liver clock, specifically in the presence of microbes results in a significant increase of abnormal connections between transcripts belonging to key metabolic pathways involved in the regulation of GNG. This demonstrates the combinatorial action between gut microbes and liver clock in the overall organization of hepatic metabolic gene transcription over time.

These data suggest both the liver clock and gut microbes aid in maintaining temporal co-expression of critical metabolic functional outputs of hepatic transcripts. Loss of either driver results in the emergence of abnormal connections, specifically those involved in carbohydrate and lipid metabolic pathways.

### Interactions between liver clock and gut microbes result in altered lipid vs carbohydrate fuel utilization

Given that loss of liver clock and gut microbes alters glucose and lipid metabolism, we interrogated how behavior, fuel utilization, and fuel switching were impacted *in vivo* via indirect calorimetry measurements over 4 days with the Promethion High-Definition Multiplexed Respirometry System. First, we detected no differences in basal metabolic rate regardless of *Bmal1* or microbial status (**Figure S6A**). While we observed no difference in overall food intake between genotypes (**Figure S6B**), we noted that SPF LKO mice exhibited slightly elevated ratio of total food intake compared to WT during the active period and less during the rest period, and no difference in GF (**Figure S6C**). Plotting hourly food intake revealed that feeding onset (indicated by arrows) was more robust in SPF LKO mice compared to WT, while no differences were evident in GF mice (**Figure S6D**). The altered feeding bouts in SPF LKO could contribute to the increased oscillating microbes over a 24-hr LD cycle, as observed in the 48-hr stool analysis (**Figure 3**). Interestingly, ambulatory motion over the same period was not different in SPF mice, however GF LKO mice exhibited increased total ambulatory motion relative to their WT counterparts (**Figure S6E,F**). This indicates a liver-clock-microbiota interaction on feeding behavior, while shifts in ambulation due to a disrupted liver clock are exaggerated by a lack of gut microbes.

We next examined energy expenditure (EE) via oxygen consumption (VO_2_) and fuel utilization via respiratory exchange ratio (RER; CO_2_ produced/O_2_ consumed). While no differences were detected between SPF genotypes during the active phase, we observed slight but significantly increased EE in SPF LKO compared to WT during the rest phase (**Figure 7A,B**). These patterns were not evident in GF conditions. We then measured RER and revealed that SPF LKO mice exhibited significantly increased RER during the active period, implying a greater utilization of carbohydrates and reduced utilization of lipids (**Figure 7C,D**). This suggests that a dysfunctional liver clock drives reduced utilization of lipids for fuel when microbes are present, supporting our evidence that GNG is also reduced. Conversely, no difference in active RER was detected in GF mice regardless of genotype (**Figure 7C,D**), which supports our evidence that LKO mice exhibit reduced GNG compared to WT in SPF, but not in GF conditions (**Figure 1E**). Interestingly, we observed decreased RER during the rest period only in GF LKO mice compared to GF WT, but not under SPF conditions (**Figure 7C,D**). This indicates that absence of both a liver clock and gut microbes may in fact enhance oxidation of lipids relative to GF WT mice.

**Figure 7.**
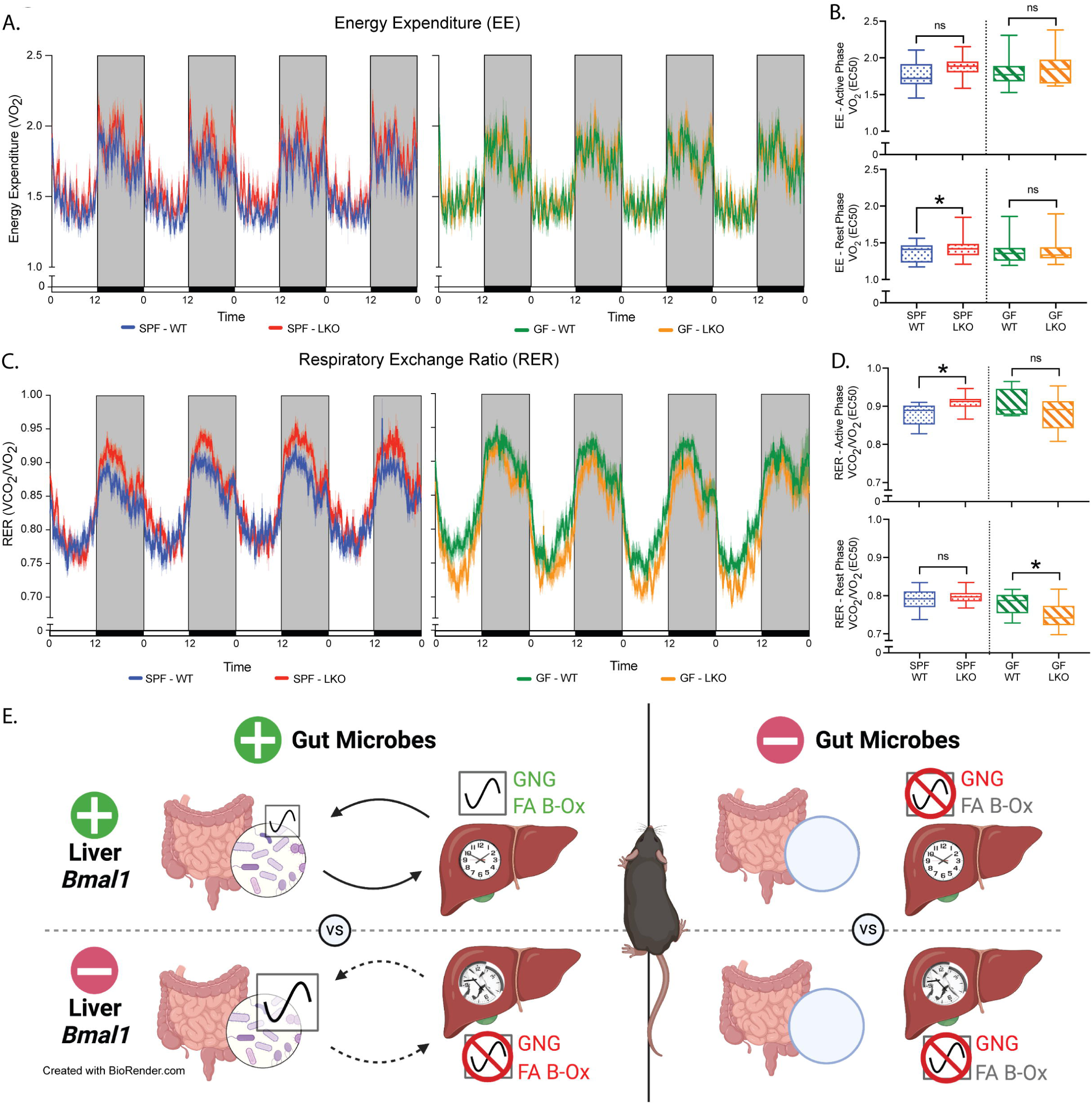
Liver clock and gut microbes differentially alter diurnal patterns of energy expenditure and fuel utilization. Indirect calorimetry assessment of SPF and GF, WT and LKO male mice, measured over 4 consecutive 12:12 LD cycles (n=12-13). **(A)** Energy expenditure (EE) represented as VO_2_. **(B)** EE divided into active (dark) and rest (light) periods, summarized by EC50 values within each period. **(C)** Respiratory Exchange Ratio (RER) represented as VCO_2_/VO_2_. **(D)** RER during active (dark) and rest (light) phases, summarized by EC50 values. Data points represent mean±SEM, box plots represent median±min/max. *p<0.05, ns=not significant. **(E)** Model figure. In SPF conditions, the liver circadian clock drives normal GNG and FA β-Ox, fecal microbial abundance oscillations, and hepatic transcriptome oscillations. Following hepatic *Bmal1* deletion, GNG and FA β-Ox are reduced, oscillating microbiota increase, and oscillating hepatic transcripts are not enriched for metabolic pathways, including GNG and FA β-Ox. In GF conditions, GNG is reduced, and oscillating hepatic transcripts are not enriched for GNG and FA β-Ox metabolic pathways regardless of liver clock functionality. Green indicates upregulation, red indicates downregulation. Solid arrows indicate intact communication, dashed arrows indicate broken communication.

In summary, we reveal microbiota-dependent and -independent effects on liver clock-mediated fuel utilization, and reduced reliance on lipids in SPF LKO that may contribute to reduced GNG and alter global metabolic homeostatic outputs.

## Discussion

Organismal-level coordination of molecular metabolism governed by the circadian clock is critical for the temporal separation of biochemical processes to maintain energy demands over a 24-hour period. Dysregulation or improper partitioning of these key processes can substantially contribute to metabolic disease development, such as obesity, non-alcoholic fatty liver disease, and type 2 diabetes. The liver is one of the most metabolically active organs, contributing to glucose and lipid homeostasis. Our data demonstrate a bidirectional and cooperative relationship between diurnal patterns of gut microbes and the liver circadian clock that aids in coordination of GNG, lipid metabolism, and fuel utilization, as outlined in **Figure 7E**.

This highly complex, coordinated, reciprocal dialogue between these two systems results in refined shifts that drive mammalian global metabolic homeostasis. We reveal that the liver clock serves as the primary driver of diurnal transcriptional networks essential for maintenance of whole-body metabolism, whereas gut microbes are a secondary, yet essential partner that translates environmental cues (e.g. what, when, and how much diet is consumed) to enhance temporal organization of clock-mediated hepatic gene expression. When either system is absent or dysfunctional, disorganization and mistiming of normally coordinated processes occurs and homeostatic mechanisms are lost. These findings provide an initial basis for the interrogations of the mechanistic underpinnings of key host-microbe circadian interactions that direct metabolism in a tissue-specific manner.

A major finding of our study is that contributions of both gut microbes and the hepatic tissue-specific circadian network mediate glucose homeostatic outcomes, which occurs independent of the central clock located within the brain (**Figure 1C,E; Figure S1**). While others have separately implicated gut microbes (Krisko et al., 2020; Wichmann et al., 2013; Zarrinpar et al., 2018) or the liver circadian clock (Lamia et al., 2008) in the regulation of GNG and glucose tolerance, respectively, we show that these systems work in conjunction to drive host glucose homeostasis. This appears to be a GNG-specific effect, supported by our evidence that both insulin sensitivity and hepatic glycogen levels were largely unaffected by LKO or GF conditions (**Figure S1C,E**). We find that gut microbes provide essential cues that mediate GNG which can only be appropriately coordinated with the liver clock, a primary driver of maintaining metabolic partitioning. This engagement in microbiota replete conditions underpins shifts between glucose and lipid metabolism for fuel utilization. By investigating these animals under GF conditions or following antibiotic-induced depletion of gut microbes in SPF mice (**Figure 2**), we were able to demonstrate that gut microbes contribute to these outcomes, serving in essence as a rheostat to impart signals in real-time that fine-tune and modulate glucose and lipid metabolism. In the presence of gut microbes, gene-targeted deletion of the primary hierarchical driver, hepatic *Bmal1,* results in reduced host catabolic processes, including GNG, that are essential to maintain metabolic homeostasis. Together, these data underscore a key role for liver clock-microbiota crosstalk in maintaining circulating glucose levels, particularly during fasting conditions when GNG is critical. Whereas each system is greatly impacted by environmental signals that influence metabolic outcomes, understanding their bidirectional dialogue is essential to co-manipulate these systems in a meaningful and effective way.

In addition to differences in glucose homeostasis determined via direct measurements of GNG, we also identified that hepatic *Bmal1* and gut microbes are integral to coordinate diurnal patterns in lipid metabolism and subsequent fuel utilization via transcript analysis and indirect calorimetry, respectively (**Figure 5, 7**). These findings, particularly in GF animals, provide additional insights into the role of liver *Bmal1* in these processes, where previous work performed by our group and others in conventionally-raised *Bmal1* WT vs. LKO mice showed the liver clock regulates lipid homeostasis via mechanisms involving AKT activation and m_6_A RNA methylation (Zhong et al., 2018; Zhang et al., 2014). Due to the significant influence that FA β-Ox imparts on GNG rates (Jones, 2016), it is possible that our observations of liver clock-mediated GNG outputs under SPF and GF conditions may be a direct result of reduced hepatic FA β-Ox flux. Previous studies showed GF mice with a functional liver clock exhibit upregulation of FA β-Ox, even under high fat diet-fed conditions (Bäckhed et al., 2007), which could be due, in part, to differential regulation of hepatic PPAR signaling (Montagner et al., 2016) and may provide protection against high-fat diet-induced obesity. Our studies in GF WT and LKO mice suggest that gut microbes also interact with the liver clock in the regulation of lipid metabolism, yet the precise microbial component driving these outcomes remains to be determined. Given the relationship between FA β-Ox and GNG, we must consider the potential effects that high-fat diet may have on the individual outputs of these pathways and their co-regulation. For instance, a previous study revealed that high fat dietary intake in *Bmal1* liver-deficient mice resulted in increased body weight gain and disrupted mitochondrial functional outputs compared to WT counterparts (Jacobi et al., 2015). We suspect that exposure to high-fat diet would impact the liver clock-microbiome relationship, perhaps exacerbating the lipid and associated GNG outputs that we observe under regular chow-fed conditions in SPF mice. This will be a focus of future studies for our group.

In addition to the metabolic implications of gut microbes on the host, we also demonstrate bidirectionality from the host liver clock to gut microbes. Here, functionality of the liver clock impacts diurnal oscillations of fecal microbiota relative abundances. Loss of a primary hepatic driver, such as *Bmal1*, results in the emergence of downstream secondary microbiota oscillators that could directly feedback onto the host, disrupting proper feedback mechanisms. Previously, Liang et al., (2015) showed that global loss of *Bmal1* in mice abolished fecal microbial abundance rhythms compared to WT. In contrast, our work shows that absence of tissue-specific *Bmal1* results in an almost 2-fold increase in oscillating ASVs, particularly within the families Lachnospiraceae and Ruminococcaceae (**Figure 3**). This indicates that the tissue-specific liver clock is a key player in maintaining normal rhythmicity of specific oscillating gut microbiota. Others have shown that these bacteria families positively correlate with and contribute to secondary bile acid metabolism (Devlin and Fischbach, 2015; Theriot et al., 2016), which are known to regulate CRs and metabolism (de Aguiar Vallim et al., 2013; Govindarajan et al., 2016). Further, previous studies have revealed that PPAR signaling is linked to bile acid synthesis and regulation (Burri et al., 2010; Li and Chiang, 2009). In fact, we observed reduced levels of transcripts involved in PPAR signaling in hepatic *Bmal1* deficient mice. Whether the observed gain in oscillating ASVs in LKO SPF mice contributes to altered circulating bile acid pools that impact glucose metabolism remains to be investigated. The concept of gain in oscillation has been previously identified in the context of microbiota relative abundances and associated metabolites in metabolic disease (Leone et al., 2015; Reitmeier et al., 2020), yet their precise meaning remains unclear. Importantly, not only do we observe a gain in oscillations of gut microbiota, but we also revealed an increase in oscillations of the host liver transcriptome in absence of either a functional liver clock or gut microbes (**Figure 5A,C**). These gains of rhythmicity confer differential enrichment of key metabolic pathways compared to SPF WT. This finding corresponds to previous observations that the emergence of unique oscillations can significantly alter functional transcriptome enrichment that impacts metabolic homeostasis (Eckel-Mahan et al., 2013; Solanas et al., 2017; Thaiss et al., 2016).

Our study underscores that gut microbiota play a key role in mammalian global metabolic homeostasis which is mediated through interactions and coordination with primary drivers of hepatic circadian clock networks. Identification of the molecular signals derived from the gut microbiome that impart liver clock-mediated GNG and overall metabolic organization is a necessary next step to gain further insights. Thus far, our results have ruled out short-chain fatty acids as a potential mediator of this interaction (data not shown). Recent studies have identified novel microbial metabolites that can be explored in our model system, and understanding how these microbial mediators interact with circadian networks particularly in human subjects will be an important step in the field. Together, our physiological and multi-omic data highlight key communication between the host hepatic circadian clock and gut microbiota, underscoring the importance of proper diurnal coordination of these two systems. These findings could have broad translational implications for how these two systems could be targeted to improve metabolic homeostasis in humans.

## Supporting information

Key Resources Table

## Acknowledgements

This work was supported by NIH/NIDDK grants R01 DK115221 (EBC), P30 DK42086 (EBC), K01 DK111785 (VAL), F31 DK122714 (KF), and T32 DK070774 (KF). This work was also supported by the University of Chicago GI Research Foundation. We thank the University of Chicago Genomics Facility (RRID:SCR_019196), the Animal Resources Center and the Gnotobiotic Research Animal Facility at the University of Chicago, and the High-Throughput Genome Analysis Core at Argonne National Laboratory. We thank Francine Vukovich for administrative support, and Dr. Matthew Brady for providing manuscript edits.

## Author Contributions

Conceptualization, K.F., V.A.L., and E.B.C.; Formal Analysis, K.F., Su.M., O.D., K.C., and V.A.L.; Investigation, K.F., Su.M., K.C., Sa.M., J.M., M.S., A.T., and V.A.L.; Resources, M.I. and J.S.T.; Writing – Original Draft, K.F., V.A.L., and E.B.C.; Writing – Review & Editing, K.F., M.C.R., V.A.L., and E.B.C.; Funding Acquisition, K.F., M.C.R., V.A.L., and E.B.C.

## Inclusion and Diversity

One or more of the authors of this paper self-identifies as a member of the LGBTQ+ community. While citing references scientifically relevant for this work, we also actively worked to promote gender balance in our reference list.

## Supplemental Tables and Figure Titles and legends

Supplementary Table S1, related to Figure 5

**Figure S1 related to Figure 1.**
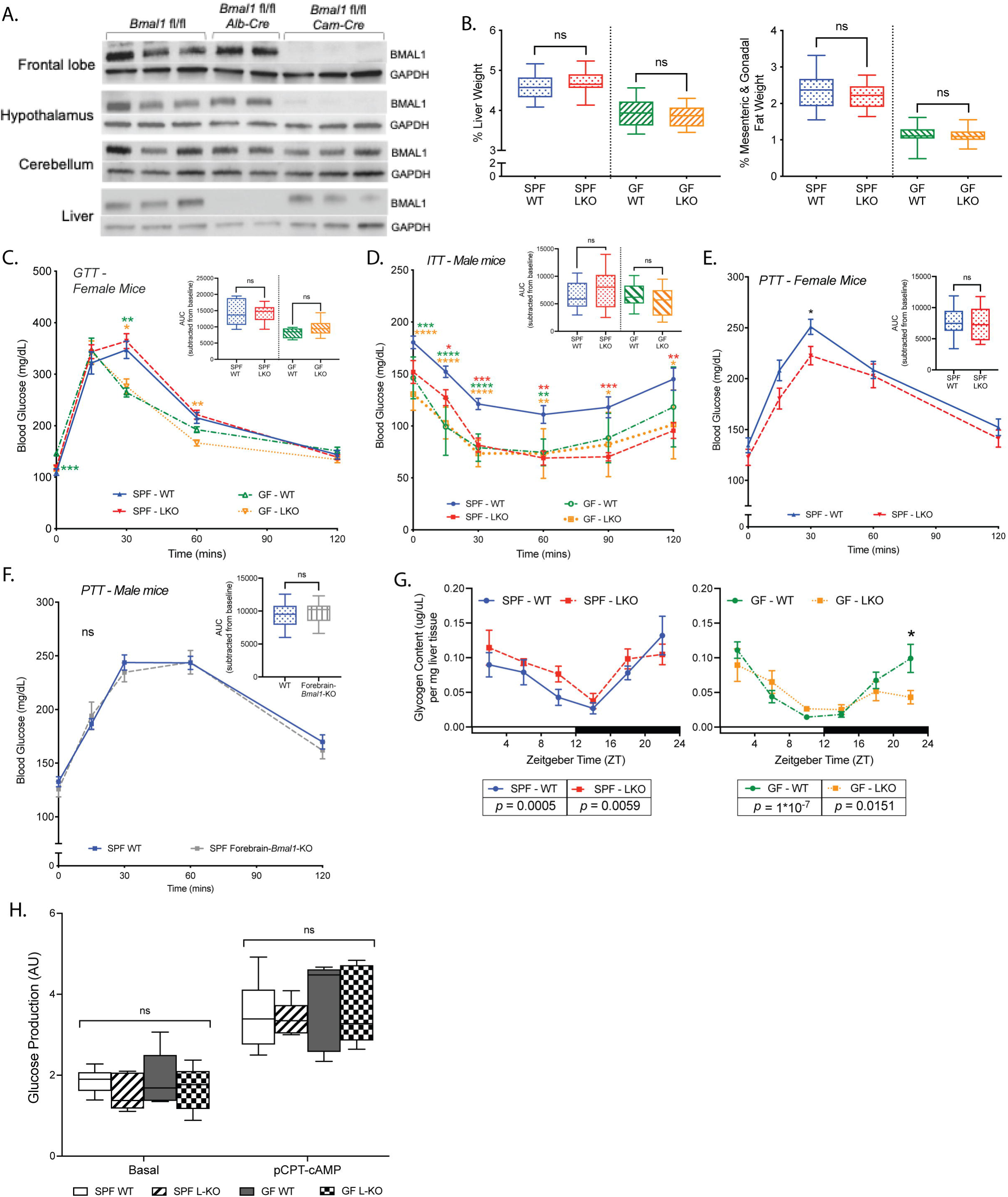
The liver circadian clock and gut microbes do not mediate insulin tolerance, hepatic glycogen, or ex vivo hepatic GNG. **(A)** Western blot of BMAL1 and GAPDH (control) in brain (frontal lobe, hypothalamus, cerebellum) and liver protein extracts collected at ZT16 from male *Bmal1^f/f^*(WT), *Bmal1^f/f^ Alb-Cre* (LKO), and *Bmal1^f/f^ Cam-Cre* (Forebrain-*Bmal1*-KO). **(B)** Liver, mesenteric & gonadal fat weight from SPF and GF, WT and LKO male mice, presented as percent of body weight (*n*=27- 34/group). **(C)** GTT in SPF and GF, WT and LKO female mice (*n*=8-10/group). **(D)** Intraperitoneal Insulin Tolerance Test in SPF and GF, WT and LKO male mice (ITT, n=11- 17/group). **(E)** PTT in SPF WT and LKO female mice (*n*=13-14/group). **(F)** PTT in SPF WT and Forebrain-*Bmal1-*KO male mice (*n*=11-16/group). **(G)** Glycogen content in liver tissue from SPF and GF WT vs LKO male mice collected every 4 hrs over 24 hrs from animals maintained in 12:12 LD (*n*=5-7/group); CircWave statistics indicate significantly oscillating (p<0.05) or not oscillating (p>0.05) values. **(H)** Glucose production by primary hepatocytes, isolated from SPF and GF WT vs LKO male mice, following 11 hrs of treatment with vehicle control (Basal) vs pCPT-cAMP, normalized to protein content and represented as arbitrary units (AU) (*n*=5/group). Data points represent mean±SEM, box plots presented as median±min/max. ****p<0.0001, ***p<0.001, **p<0.01, *p<0.05, ns=not significant; colored stars represent significance relative to SPF WT. Inset graph represents AUC normalized to baseline glucose values.

**Figure S2 related to Figure 2.**
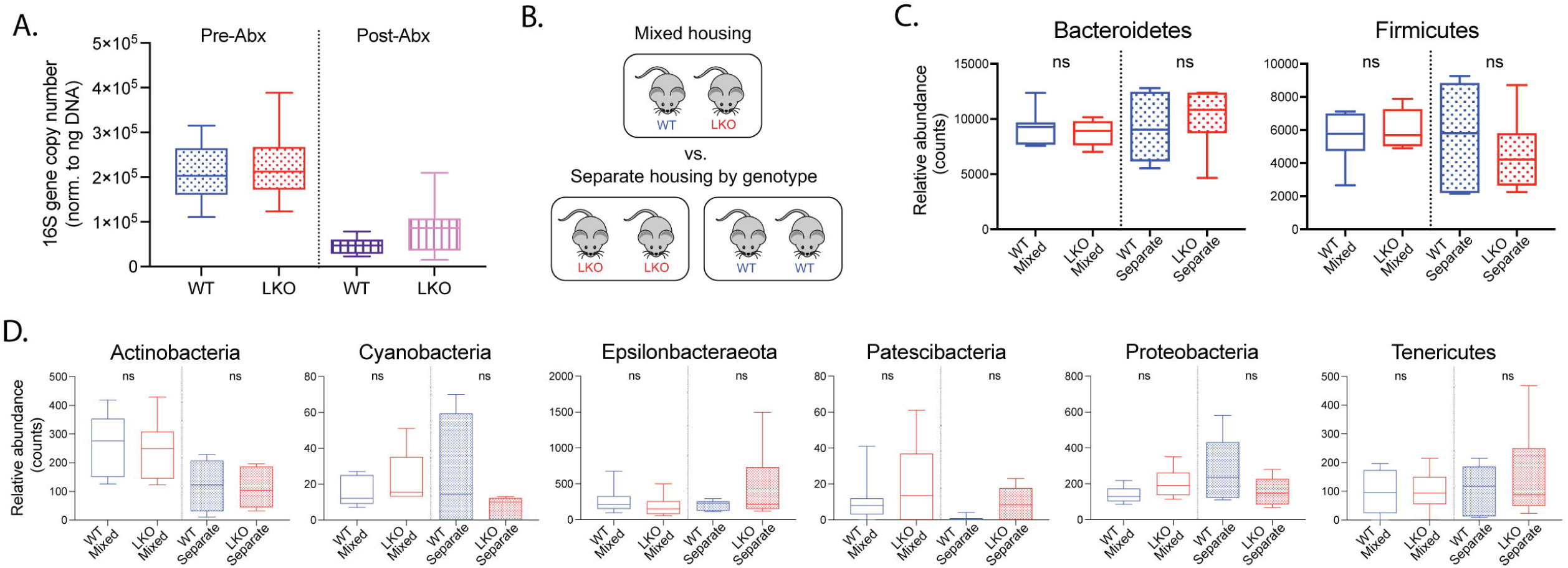
Liver circadian clock status does not impact gut microbial community membership. **(A)** 16S rRNA gene abundance in stool collected before (Pre-Abx) and after (Post-Abx) antibiotic treatment, measured via quantitative PCR and normalized to DNA concentration. **(B)** Experimental scheme of mixed vs. separate housing at time of weaning. **(C,D)** 16S rRNA gene sequencing analysis of stool collected from WT vs LKO male mice housed either by mixed genotypes (WT+LKO) or via separate housing of genotypes (WT+WT and LKO+LKO) (*n*=6- 7/group); relative abundance counts of major (C) and minor (D) phyla. Box plots represent median±min/max. ns=not significant.

**Figure S3 related to Figure 3.**
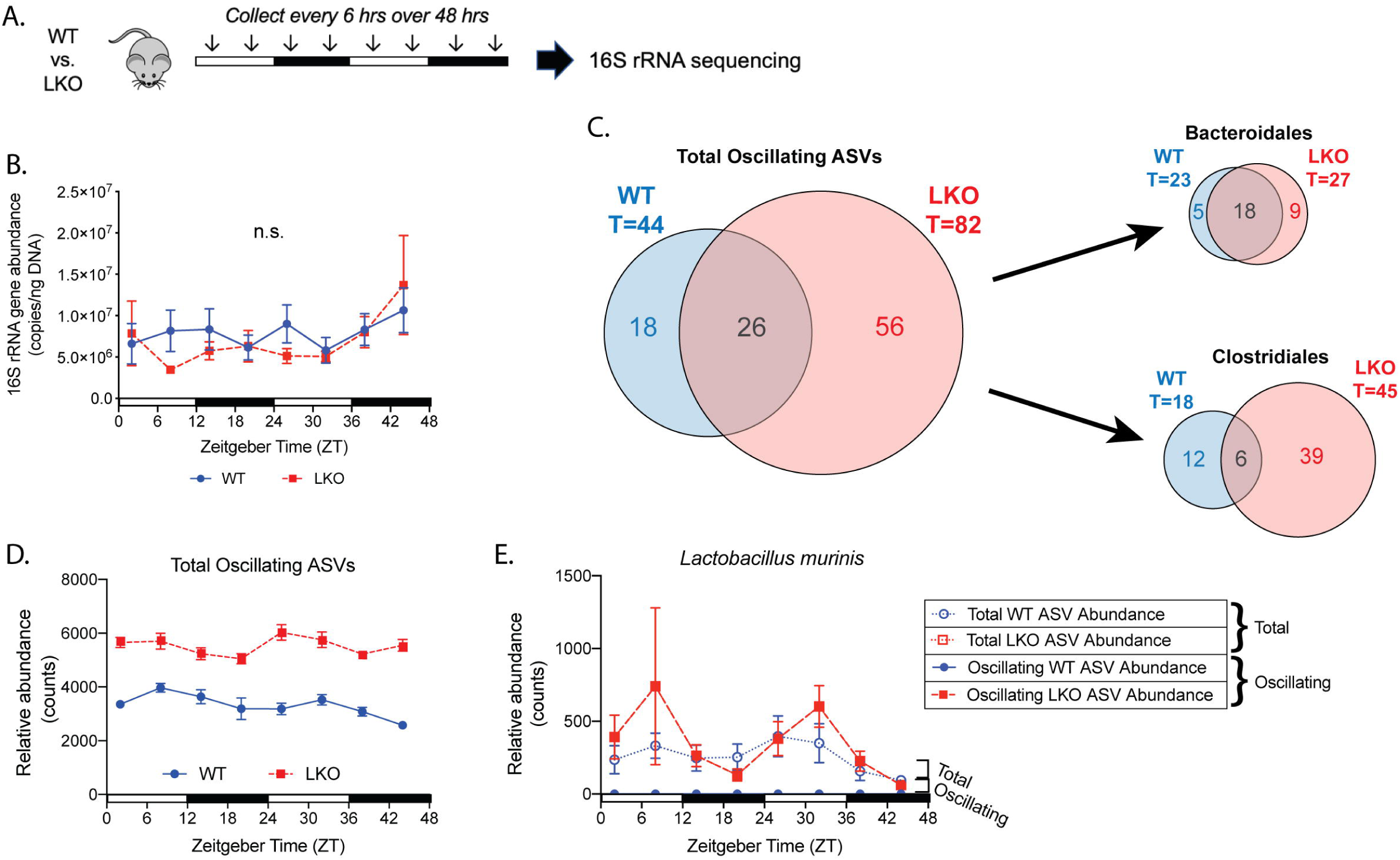
Loss of a functional liver clock results in increased abundance of oscillating ASVs. 16S rRNA gene sequencing analysis of stool sampled from WT and LKO male mice every 6 hrs over two consecutive 24-hr LD cycles (8 samples per animal) via repeat collections from the same mouse (*n*=7-8/group). **(A)** Experimental scheme of 48-hr stool collection. **(B)** 16S rRNA gene abundance measured via qPCR normalized to DNA concentration. **(C)** Venn diagrams presenting total, shared, and unique oscillating ASVs among all taxonomic groups, as well as within specific classes Bacteroidales and Clostridiales. **(D)** Relative abundance counts of all oscillating ASVs. **(E)** Relative abundance counts of total vs oscillating ASVs within *Lactobacillus murinus*. Data points represent mean±SEM. ns=not significant.

**Figure S4 related to Figure 4.**
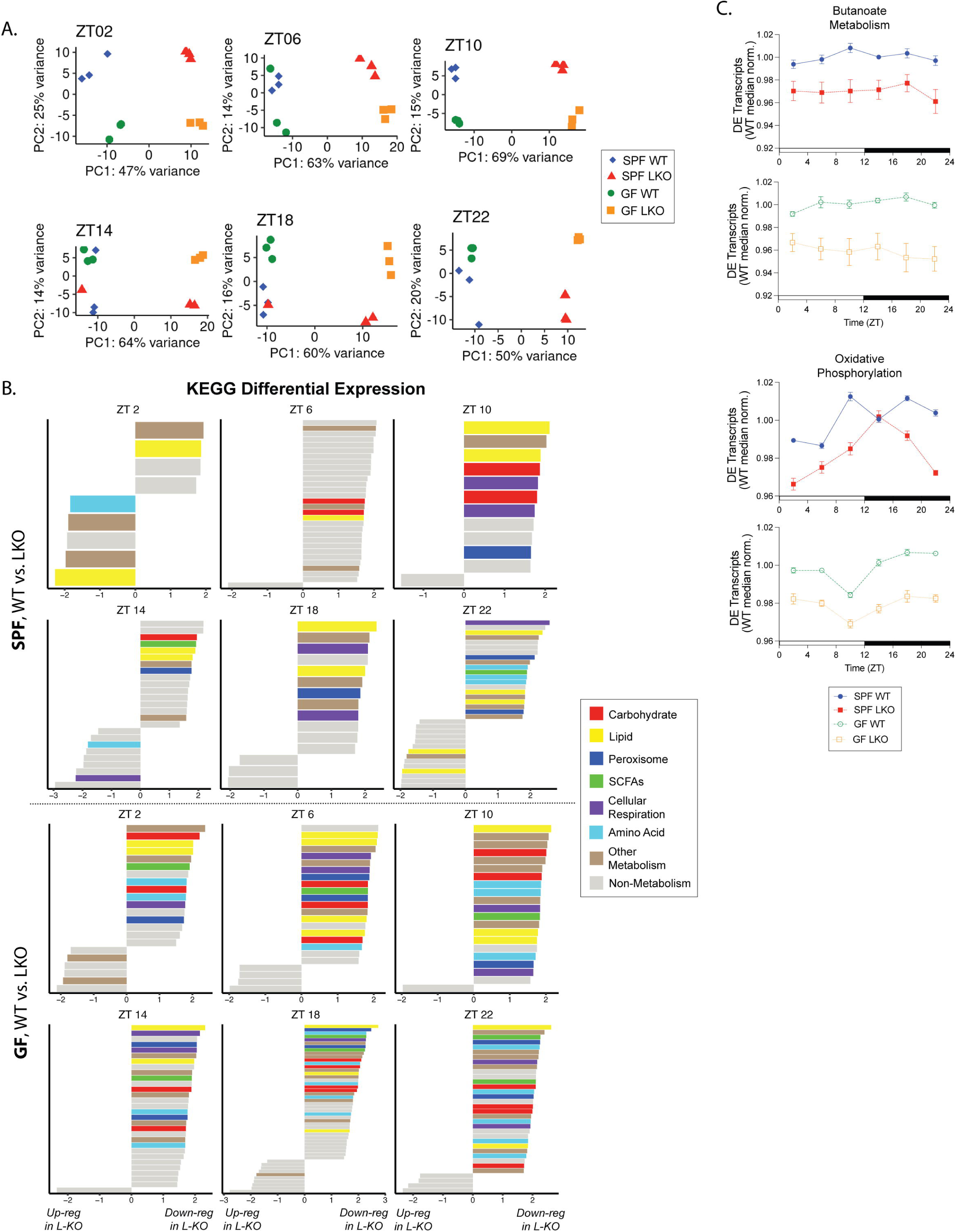
Liver-clock downregulation of metabolic pathways is evident across all timepoints. Differential transcriptome analysis of liver samples collected every 4 hrs over 24-hrs from animals maintained in 12:12 LD (ZT2, 6, 10, 14, 18, 22) from SPF and GF, WT and LKO male mice (n=3/timepoint/group). **(A)** Principal Component Analysis of transcriptome profiles, samples divided by collection timepoint. **(B)** Differentially regulated KEGG pathways between WT and LKO, within SPF and GF, divided by ZT collection timepoint. Metabolic pathways are highlighted according to the legend, and non-metabolic pathways are colored gray. **(C)** WT- median-normalized expression of differentially expressed (DE) genes within identified KEGG pathways.

**Figure S5 related to Figure 5 & 6.**
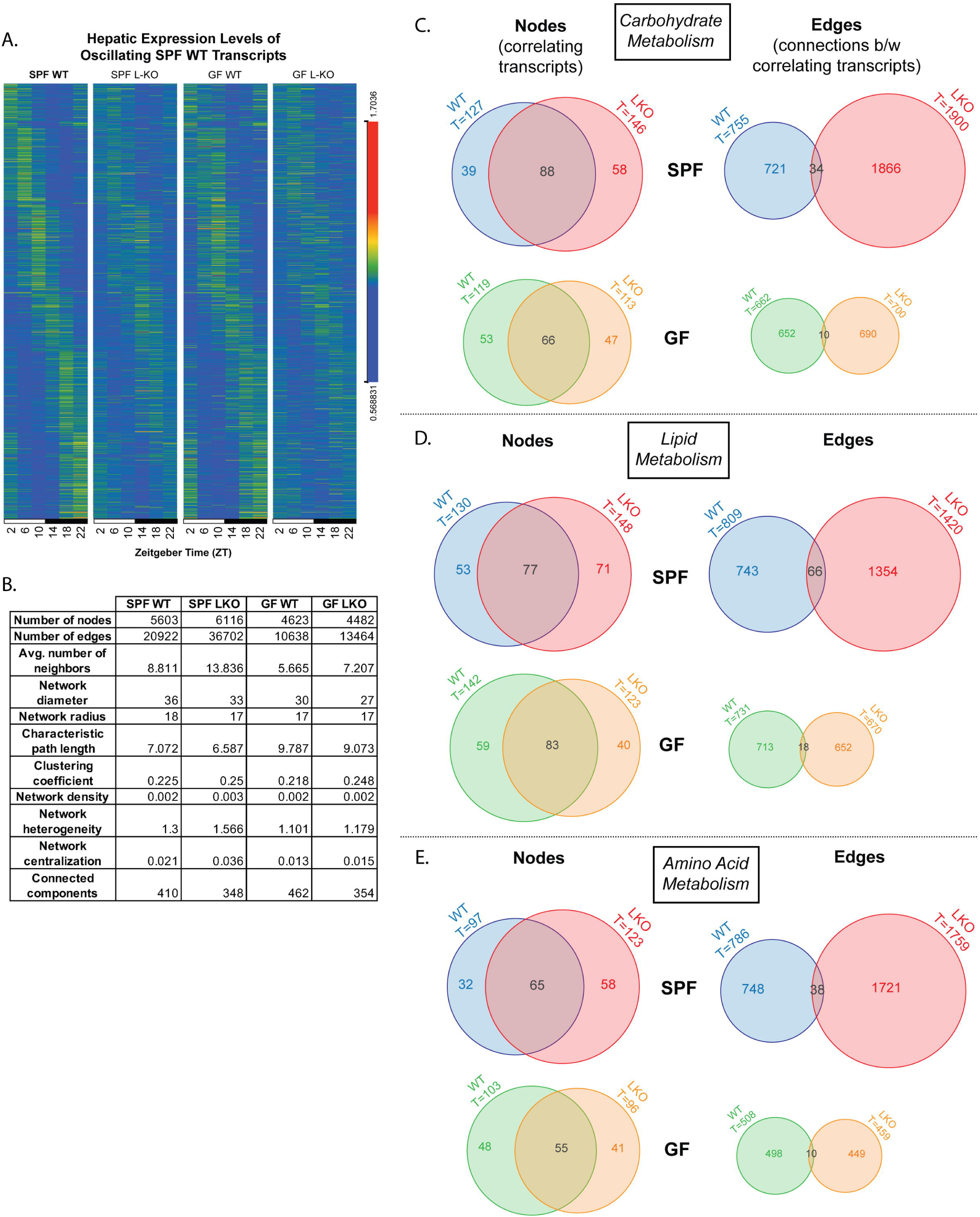
Liver clock drives increases in unique hepatic transcript oscillations and co-occurrence expression in the presence of gut microbes. Diurnal and network transcriptome analysis of liver samples collected every 4 hrs over 24-hrs from animals maintained in 12:12 LD (ZT2, 6, 10, 14, 18, 22) from SPF and GF, WT and LKO male mice (n=3/timepoint/group). **(A)** Transcript expression profiles of significantly oscillating SPF WT transcripts; expression values normalized by median, transcripts ordered by time of maximum expression and phase. **(B)** Network co-occurrence statistics. **(C-E)** Venn diagrams summarizing correlating transcripts (nodes) and connections (edges) that annotate to carbohydrate **(C)**, lipid **(D)**, and amino acid **(E)** metabolic pathways by KEGG, and crossover within SPF and GF conditions; network co-occurrence analysis of correlating transcripts over time within each group (p<0.001).

**Figure S6 related to Figure 7.**
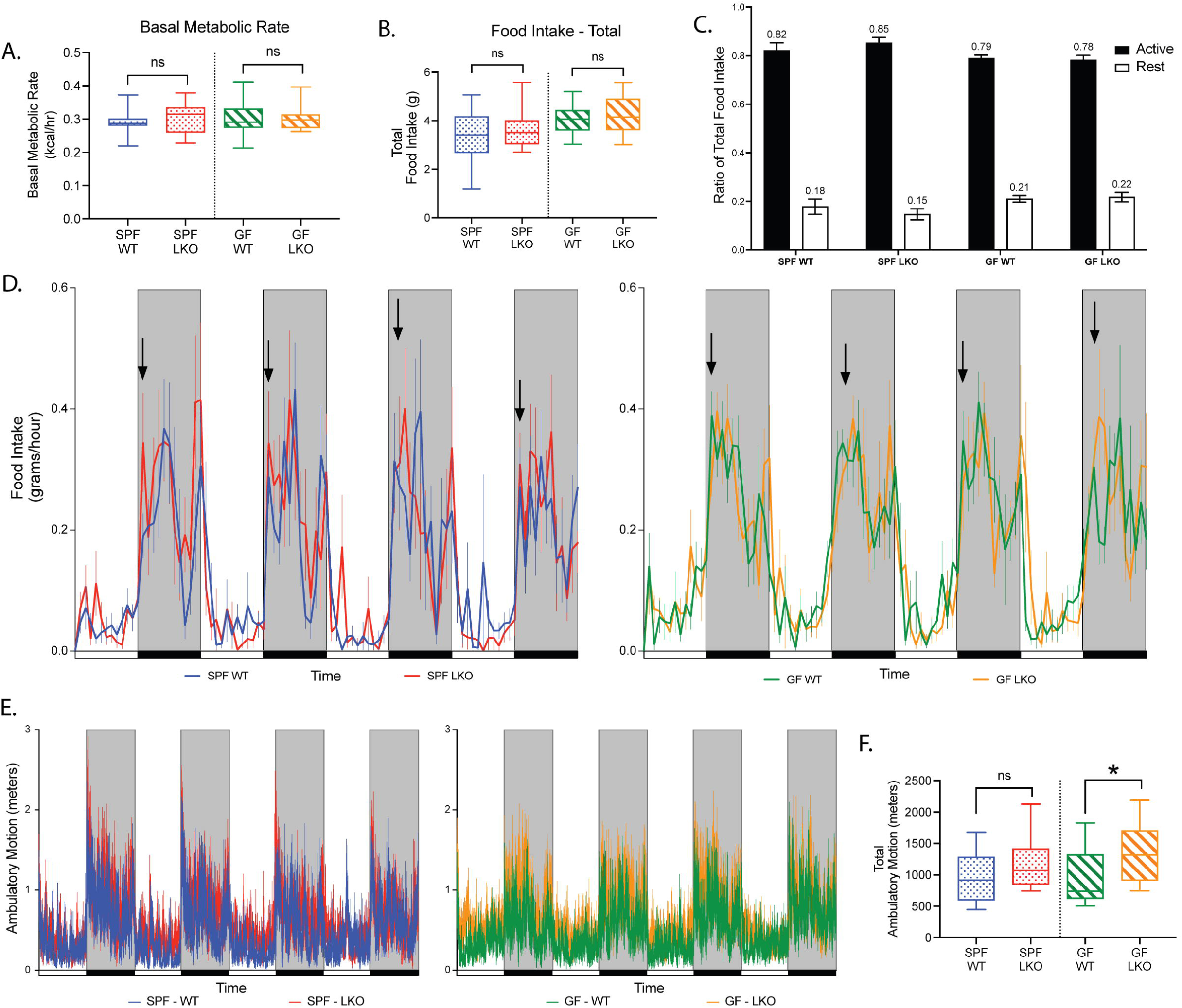
Liver clock and gut microbes convey modest changes in food intake patterns and locomotor activity. Indirect calorimetry assessment of SPF and GF, WT and LKO male mice, measured over 4 consecutive 12:12 LD cycles (n=12-13). **(A)** Basal metabolic rate represented as kcal/hr. **(B)** Total grams of food intake per LD cycle. **(C)** Ratio of total food intake consumed during the active vs rest periods. **(D)** Food intake summarized as grams consumed per hour; arrows indicate time of feeding onset during the active period. **(E)** Ambulatory motion measured by meters. **(F)** Total ambulatory motion measured by meters per LD cycle. Data points and bars represent mean±SEM, box plots represent median±min/max. *p<0.05, ns=not significant.

## STAR Methods

### Contact for Reagent and Resource Sharing

#### Lead contact

Further information and requests for resources and reagents should be directed to and will be fulfilled by the Lead Contacts, Eugene Chang and Vanessa Leone.

#### Materials Availability

This paper does not report original reagents or materials.

#### Data and Code Availability

16S rRNA amplicon raw sequencing files are deposited in the NCBI Sequence Read Archive. RNA sequencing files are available for download at the NCBI Gene Expression Omnibus. Accession numbers are listed in the key resources table.

This paper does not report original code.

Any additional information required to reanalyze the data reported in this paper is available from the lead contact upon request.

### Experimental Model and Subject Details

#### Animals

All animal protocols and experimental procedures were approved by the University of Chicago Institutional Animal Care and Use Committee (IACUC). All mice used in these studies were on a C57BL/6J background. Specific pathogen-free (SPF) *Bmal1^fl/fl^*and Albumin-Cre male and female mice were purchased from the Jackson Laboratory and bred in the UChicago vivarium as described in Lamia *et al*. (2008). SPF *Bmal1^fl/fl^* Alb-Cre mice were re-derived under germ-free conditions and maintained in flexible film isolators (CBC Ltd. Madison, WI, USA) in the UChicago Gnotobiotic Research Animal Facility. SPF *Bmal1^fl/fl^*mice were crossed with *CamiCre* mice (a generous gift from Dr. Joseph Takahashi, as described in Izumo et al., 2014) to achieve forebrain-specific knockout of *Bmal1*. All mice were maintained under standard 12:12 light/dark conditions (lights on beginning at 6am, Zeitgeber (ZT) 0) and fed autoclaved *ad libitum* JL Rat and Mouse/Auto 6F 5k67 (LabDiet, St. Louis, MO, USA) for at least 2 weeks prior to and throughout all experiments. Body weight and food consumption were monitored weekly throughout the studies. At 10-14 weeks of age, mice were acclimatized to individual housing for 14 days, after which stool was collected every 6 hours for 48 hours. After 3 weeks, mice were sacrificed via CO_2_ asphyxiation followed by cervical dislocation over a 24-hour period at six ZT time points: ZT2=8 AM, ZT6=12 PM, ZT10=4 PM, ZT14=8 PM, ZT18=12 AM, and ZT22=4 AM.

Immediately prior to sacrifice, basal blood glucose was measured from tail snip blood via Accu- Check Compact Plus Diabetes Monitoring Kit and test strips (Roche Diagnostics, Indianapolis, IN, USA), or OneTouch Ultra 2 Blood Glucose Monitoring System (OneTouch, LifeScan, Malvern, PA, USA). Blood was also collected via heparin-coated microvette tubes (Sarstedt, Numbrecht, Germany) for insulin measurement by the Ultra-sensitive Insulin ELISA (ALPCO, Salem, NH, USA). Liver, plasma, brain, and intestinal luminal contents were snap frozen in liquid nitrogen and stored at -80C.

#### Conventionalization Studies

For conventionalization, fresh stool from adult C57BL/6J male mice was collected and immediately resuspended in sterile PBS (100mg stool/1mL PBS) and vortexed, followed by a brief spin to remove debris. Individually housed GF *Bmal1^fl/fl^*or *Bmal1^fl/fl^-AlbCre* 11- to 13-week-old *ad libitum*-fed male mice were orally gavaged with 150 uL of this suspension. Body weights and food consumption were monitored weekly.

#### Antibiotic-treatment Studies

For antibiotic treatment, SPF *Bmal1^fl/fl^*or *Bmal1^fl/fl^-AlbCre* 11- to 13-week-old male mice were administered an antibiotic cocktail consisting of vancomycin (0.5 mg/mL), neomycin (1 mg/mL), and cefoperazone (0.5 mg/mL) prepared in autoclaved water and sterile-filtered. The protocol involved using a combination of daily gavage for 7 days followed by incorporation and *ad libitum* delivery in drinking water for 1 additional week. Body weights were monitored daily.

#### Primary hepatocyte Isolation & Culture

Isolation and culture of primary hepatocytes was performed as described in http://mouselivercells.com. 15-week-old SPF and GF *Bmal1^fl/fl^* and *Bmal1^fl/fl^-AlbCre* male mice were anesthetized (90 mg ketamine/kg body weight, 6 mg xylazine/kg body weight), perfused via portal vein cannulation with Collagenase Type IV (Worthington Biochemical) *in situ*, digested in low-glucose DMEM (Corning), and isolated in high-glucose DMEM (Gibco). Live hepatocytes, determined via Trypan blue exclusion, were seeded at a density of 600,000 hepatocytes in 6- well collagen-coated plates (Thermo Scientific) and cultured in low-glucose DMEM (Corning) for two days with daily media change prior to initiating downstream assays.

### Method Details

#### Oral Glucose, Intraperitoneal Pyruvate, and Intraperitoneal Insulin Tolerance Tests

For glucose (GTT) and pyruvate (PTT) tolerance tests, 12- to 16-week-old male GF and SPF mice were fasted overnight for 14 hours, starting at ZT12. For insulin tests (ITT), mice were fasted for 5 hours, starting at ZT22. Baseline blood glucose was measured via tail snip using a hand-held glucometer. Mice were administered either an oral bolus of 20% dextrose (Hospira) in sterile water solution (2g/kg body weight), an intraperitoneal (IP) injection of sterile-filtered sodium pyruvate (Sigma-Aldrich) in PBS (2g/kg body weight), or an IP injection of 0.1U/mL Humulin R Insulin (Eli Lilly) in PBS (1U/kg body weight). For GTT and PTT, blood glucose was then measured at 15-, 30-, 60-, and 120-minutes post gavage or injection. For ITT, blood glucose was then measured at 15-, 30-, 60-, 90-, and 120-minutes post injection. During GTT, insulin levels were determined at baseline, 30, 60, and 120 minutes in blood collected in heparin-coated microvette tubes (Sarstedt, Numbrecht, Germany) using the Ultra-sensitive Insulin ELISA (ALPCO, Salem, NH, USA). Area under the curve (AUC) was calculated and compared between genotypes using GraphPad Prism v9.

#### Western Blot

Liver and brain tissue (5-10 mg) were collected at ZT16 and placed in 250-500 µL of ice-cold RIPA lysis buffer (Thermo Scientific RIPA Lysis and Extraction Buffer, cOmplete Mini Protease Inhibitor Cocktail, 100µM PMSF). Protein concentrations were determined via Bicinchoninic Acid (BCA) Protein Assay (Thermo Scientific) and 30µg of protein was separated on 4-20% Mini-PROTEAN gel (Bio-Rad) and transferred to a PVDF membrane (Millipore). Membranes were blocked with 5% nonfat milk in Tris-buffered saline (TBS) (20 mM Tris pH7.6, 150 mM NaCl) and incubated overnight at 4°C in 5% nonfat milk in TBS-Tween (TBS-T) (20 mM Tris pH7.6, 150 mM NaCl, 0.1% Tween-20) containing primary antibodies: anti-BMAL1 (1:1000; Abcam) or anti-GAPDH (1:100000; Invitrogen). Membranes were washed 6 times for 5 min in TBS-T and incubated for 1 hour at room temperature in TBS-T containing 5% nonfat milk and either anti-mouse or anti-rabbit secondary antibody (1:10,000; Abcam). Membranes were washed 6 times for 5 min each in TBS-T, developed with SuperSignal West Pico PLUS Chemiluminescent Substrate (Thermo Scientific), and exposed using the Chemi-Doc MP Imaging System (Bio-Rad Laboratories, Hercules, CA, USA).

#### Primary Hepatocyte Glucose Production Assay

Cultured mouse hepatocytes were serum-starved overnight and washed with PBS with MgCl_2_ and CaCl_2_ (Sigma-Aldrich). Hepatocytes were exposed to glucose-free DMEM (Sigma-Aldrich) (20 mM sodium lactate, 2 mM sodium pyruvate, 2 mM L-glutamine, 15 mM HEPES) with or without 0.1 mM pCPT-cAMP (Sigma-Aldrich) for 11 hours. Media was collected and glucose concentrations measured via Autokit Glucose enzymatic assay (FUJIFILM Medical Systems Inc, Lexington, MA, USA) and normalized to protein content determined via BCA assay.

#### Glycogen Assay

Frozen liver samples were weighed, and glycogen measurement was performed using the Glycogen Assay Kit II (Colorimetric) (Abcam, Cambridge, UK) following the published kit instructions.

#### Bacterial Gene Quantification

16S gene copy number was determined from stool as previously described (Leone et al., 2015; Louis et al., 2010; Vital et al., 2013). The 16S gene was quantified using a standard curve for gene copy number using primer sequences cloned into a PCR4-TOPO plasmid (see Key Resources Table for primer sequences).

#### 16S DNA Extraction, Sequencing, and Analysis

Stool was beadbeaten using 0.1-mm-diameter zirconia/silica beads (BioSpec Products, Bartlesville, OK, USA) as previously described (Leone et al., 2015). Supernatants were extracted using equal volumes of Phenol:Chloroform:Isoamylalcohol (25:24:1; Ambion, Austin, TX, USA) and DNA precipitated using an equal volume of 100% isopropanol. DNA concentration was measured using the Quant-iT PicoGreen dsDNA Assay kit (Invitrogen, Grand Island, NY) and diluted to 1-20 ng/μL. The V4-V5 region of the 16S rRNA encoding gene was amplified using standard Earth Microbiome Project protocols. Sequencing was performed at the High-Throughput Genome Analysis Core (HGAC; part of the Institute for Genomics & Systems Biology [IGSB]) at Argonne National Laboratory. Paired-end reads were imported into Quantitative Insights Into Microbial Ecology version 2 (QIIME2) software suite (Hall and Beiko, 2018) (https://qiime2.org) and demultiplexed using Dada2 (Callahan et al., 2016). To maximize sampling depth while prioritizing equal retention of samples across groups, 10598 (Figure 3 and S3, 48-hr fecal collection) or 15578 (Figure S2, co-housing vs separate housing fecal collection) reads were included per sample, determined by the mean subtracted by 1.5 times standard deviation of sample read counts across all samples within experiment. Taxonomy was compiled using the classify-sklearn plugin with Silva database version 132 2020.8 (Bokulich et al., 2018; Yarza et al., 2014).

#### Liver RNA Extraction, Sequencing, and Analysis

Total RNA was isolated by homogenizing bulk liver tissue in TRIzol (Ambion, Hampton, NH) followed by chloroform extraction, as previously described (Leone et al., 2015). RNA quality and quantity was assessed using the Agilent bio-analyzer and RNA-SEQ libraries were generated using the Illumina Stranded mRNA Prep at the UChicago Genomics Core Facility. Sequencing of mRNA directional, single-end 50 base pair reads was performed on the HiSEQ4000 with Illumina reagents and protocols. Data was demultiplexed using the Illumina bcl2fastq software. All 72 samples were run on 6 lanes, and fastq files were concatenated. Quality control was performed using FastQC v0.11.5 and MultiQC v1.10.1. Sequence alignment was performed by STAR version 2.6.1b by mapping to the mm10 whole genome (Dobin et al., 2013). Gene transcript counts were determined by Subread:featurecounts v2.0.0.

#### Differential gene expression analysis

Differential analysis was performed via DESeq2, both on all timepoints pooled and within each individual timepoint. Protein coding genes were identified via biomaRt (version 2.40.5) using GRCm39 mouse genes. Two-way analysis for each pair of experimental groups were performed. The Wald statistic, calculated via DESeq2, was used to build a ranked list of genes for each comparison for each timepoint. Fast Gene Set Enrichment Analysis (fGSEA, version 1.10.1) was utilized to identify differentially regulated pathways (with parameter nperm=10000) within multiple annotated gene sets from Molecular Signatures Database (MSigDB, version 7.0). Significantly identified pathways were filtered by adjusted p value < 0.05, and subsequently binned into categories via custom R script. fGSEA was utilized to calculated normalized enrichment score (NES) and identify leading edge genes.

#### Oscillation analysis

Feature counts were normalized by variance-stabilizing transformation (VST) in DESeq2 (version 1.24.0). VST-normalized data was used for principal component analysis (PCA). VST-normalized data was analyzed to identify significantly oscillating transcripts via empirical JTK_CYCLE (Hutchison et al., 2015). Heatmaps were generated from VST-normalized data, ordered by time of peak expression, and normalized by the median value for each transcript within each group over time, and visualized using Orange3 (https://orangedatamining.com). Metascape was utilized to identify statistically enriched pathways from each list of oscillating transcripts (Zhou et al., 2019) (http://metascape.org), which were then binned into categories via a custom R script.

#### Network co-occurrence analysis

Network co-occurrence analysis was performed using RNA-seq raw transcript abundance counts. Transcripts with low abundance counts (<50 counts in all samples or genes with 0 counts in more than 2 samples per timepoint) were removed. Spearman correlation values for comparisons between pairs of transcripts were calculated within each genotype. Transcript count values were randomly selected from one of the three samples for a given timepoint (ZT 2, 6, 10, 14, 18, 22), then correlation *p* and *r* values between transcripts over time were calculated in R, repeated over 500 permutations (*‘dplyr’, ‘rcorr’, ‘Hmisc’*). Correlation matrices for the 500 permutations were averaged to generate a single matrix of transcript correlation values. The matrix was flattened into a network-importable table format in R (*‘cormat’, ‘pmat’*): flatCorrMatrix <- function(cormat, pmat) { ut <- upper.tri(cormat) data.frame(row = rownames(cormat)[row(cormat)[ut]], column = rownames(cormat)[col(cormat)[ut]], cor =(cormat)[ut], p = pmat[ut]). Correlations were filtered for p<0.001 and spearman r>0.95, and all self-relationships removed. This filtered correlation table was used for network visualization in Cytoscape (Cline et al., 2007) (https://cytoscape.org), as well as for downstream analyses of node connectivity and centrality. Annotations from the Kyoto Encyclopedia of Genes and Genomes (KEGG) were added (ensemble.org) and utilized to identify transcript nodes in each network with specific functionally related pathways.

#### Metabolic Cage Studies

Mice were acclimatized to individual housing for 5 days prior to metabolic monitoring, which was conducted using the Promethion Metabolic System (Promethion High-Definition Multiplexed Respirometry System for Mice; Sable Systems International, Las Vegas, NV, USA). All measurements were recorded at 3 minute intervals across all cages. Food and water intake were recorded by gravimetric measurement. Physical movement was determined by infrared sensor beam breaks. Oxygen consumption, carbon dioxide production, and respiratory quotient were measured by indirect calorimetry. Energy expenditure (kcal/hr) was calculated by the Weir equation (Weir, 1949).

Basal metabolic rate (BMR) was measured on the first day of metabolic cage housing. Following transfer into metabolic cages at ZT3, food was immediately removed. During the 3-hr period between ZT7 to ZT10, the lowest average post-absorptive energy expenditure value over a 5-minute period was identified as BMR (Speakman, 2013). Food was returned at ZT10 and mice were allowed to acclimatize for 2 days, followed by 4 days of resting metabolic rate (RMR) measurements. At the end of each experiment involving GF mice, validation of GF status was performed using 16S rRNA gene PCR on DNA extracted from fecal samples collected pre- and post-metabolic caging. Any mice that were identified as positive were excluded from the study.

All data was recorded via IM3 software v21.0.2 and converted to xml via a custom Sable Systems macro. Data was wrangled and cleaned in Python, including identification of each diurnal cycle using environmental lux values. Within each time period, sum was calculated for beam breaks and movement, range was calculated for food and water intake. EC50 (Riachi et al., 2011) was calculated for RQ, VO2, and energy expenditure in R using nplr v.1-7. ANCOVA (body weight as covariant) was performed for each measurement between two groups using R.

### Quantification and Statistical Analysis

Data are presented as mean ± SEM, or box-and-whisker plots represented as median ± min/max. Statistical analysis was performed using either GraphPad Prism v9 software or R packages. Two-tailed unpaired Welch’s t tests or ANCOVA were performed between two groups, and Brown-Forsythe & Welch ANOVA followed by Dunnett’s tests were performed between three or more groups; p < 0.05 was considered statistically significant. Metascape was used to identify significantly enriched pathways from oscillation transcriptome profiles by q-value < 0.05. Spearman correlation was performed to identify significantly correlated gene expression over time by p < 0.001 and r < 0.95. fGSEA was used to identify significantly enriched pathways from differential gene expression analysis by padj < 0.05 and nperm = 10000. CircWave V1.4 (http://clocktool.org) or empirical JTK_CYCLE (Hutchison et al., 2015) were used to identify significantly oscillating data by p < 0.05 (CircWave) or GammaBH < 0.05 (p-value calculated from Gamma fit of empirical null distribution, adjusted for Benjamini-Hochberg false-discovery rate, eJTK). Statistical outliers were identified via 2 standard deviations +/- mean and removed from the analysis. Metabolic cage statistical outliers were removed based on S_n_ values (Rousseeuw and Croux, 1993) using the R package robustbase (https://robustbase.r-forge.r-project.org). A consistency factor of 1.1926 was used to calculate the S_n_ value for each channel within each cycle, and outliers were defined as +/- 3 times the S_n_ value from the median. Python v3.7.5 and R v3.6.3 were used.

### Data and Software Availability

16S rRNA sequences are available for download at the NCBI Sequence Read Archive, identified by accession PRJNA815335. RNA sequences are available for download at the NCBI Gene Expression Omnibus, identified by accession GSE184303.

