## Supplementary material for "Gut Microbes and the Liver Circadian Clock Partition Glucose and Lipid Metabolism": Key Resources Table

| REAGENT or RESOURCE | SOURCE | IDENTIFIER |
| --- | --- | --- |
| Antibodies | | |
| Rb pAb to BMAL1 | Abcam | Cat#ab93806  RRID#AB_10675117  Lot#GR3368431-1 |
| Anti-GAPDH, MAb | Invitrogen | Cat#AM4300  RRID#AB_437392  Lot#00959883 |
| Anti-mouse IgG, HRP-linked Antibody | Cell Signalling Technology | Cat#7076S  RRID#AB_330924  Lot#33 |
| Anti-rabbit IgG, HRP-linked Antibody | Cell Signalling Technology | Cat#7074S  RRID#AB_2099233  Lot#29 |
| Bacterial and Virus Strains | | |
| n/a |  |  |
| Biological Samples | | |
| n/a |  |  |
| Chemicals, Peptides, and Recombinant Proteins | | |
| 50% Dextrose Injection, USP | Hospira | Cat#00409-4902-34 |
| Sodium Pyruvate | Sigma-Aldrich | Cat#P5280 |
| Humulin R Insulin | Eli Lilly | Cat#0002-8215-01 |
| pCPT-cAMP | Sigma-Aldrich | Cat#C3912 |
| DMEM low glucose, primary hepatocyte digestion & culture medium | Corning | Cat#10-014-CV |
| DMEM high glucose, primary hepatocyte isolation medium | Gibco | Cat#11960-044 |
| Collagenase Type IV | Worthington Biochemical | Cat#LS004188 |
| PBS with MgCl_2_ and CaCl_2_ | Sigma-Aldrich | Cat#D8662 |
| HEPES buffer | Thermo Scientific | Cat#15630080 |
| DMEM without glucose, L-glutamine, phenol red, sodium pyruvate, sodium bicarbonate | Sigma-Aldrich | Cat#D5030 |
| SuperSignal West Pico PLUS Chemiluminescent Substrate | Thermo Scientific | Cat#34580 |
| iTaq Universal SYBR Green Supermix | Bio-Rad | Cat#1725124 |
| TRIzol Reagent | Ambion | Cat#1559018 |
| Phenol:Chloroform:Isoamylalcohol, 25:24:1 | Ambion | Cat#AM9732 |
| RIPA Buffer | Thermo Scientific | Cat#89900 |
| cOmplete Mini Protease Inhibitor Cocktail | Sigma-Aldrich | Cat#11836153001 |
| Vancomycin Hydrochloride from Streptomyces Orientalis | Sigma-Aldrich | Cat#V2002 |
| Neomycin, Sulfate | Fisher Scientific | Cat#BP2669-25 |
| Cefoperazone Sodium Salt | Sigma-Aldrich | Cat#C4292 |
| Ketathesia, 100 mg/mL | Henry Schein | Cat#11695-0702-1 |
| Xylazine Injection, 20 mg/mL | Akorn Animal Health | Cat#59399-110-20 |
| 2-Metcaptoethanol BP176-100 Electrophoresis | Fisher Scientific | CAS#60-24-2 |
| Bovine Serum Albumin | Sigma-Aldrich | Cat#A7906 |
| Critical Commercial Assays | | |
| Quant-iT PicoGreen dsDNA Assay kit | Invitrogen | REF#P7589 |
| Glycogen Assay Kit II (Colorimetric) | Abcam | Cat#ab169558 |
| Ultra-sensitive Insulin ELISA | ALPCO | Cat#80-INSMSU-E01 |
| Bicinchoninic Acid (BCA) Protein Assay | Thermo Scientific | Cat#PI23228 |
| Autokit Glucose | FUJIFILM Medical Systems USA | Cat#99703001 |
| Deposited Data | | |
| 16S rRNA amplicon raw sequencing reads | NCBI BioProject | <https://www.ncbi.nlm.nih.gov/bioproject/>; Accession#PRJNA815335 |
| mRNA raw sequencing reads | NCBI GEO | <https://www.ncbi.nlm.nih.gov/geo>; Accession#GSE184303 |
| Experimental Models: Cell Lines | | |
| n/a |  |  |
| Experimental Models: Organisms/Strains | | |
| Mouse: *Bmal1^flox/flox^* (B6.129S4(Cg)-*Arntl^tm1Weit^/*J) | The Jackson Laboratory | Strain#007668  RRID#IMSR_JAX:007668 |
| Mouse: Albumin-Cre (B6.Cg-*Speer6-ps1^Tg(Alb-cre)21Mgn^*/J | The Jackson Laboratory | Strain#003574  RRID#IMSR_JAX:003574 |
| Mouse: Cami-Cre (B6 background) | Dr. Joseph Takahashi | n/a |
| Oligonucleotides | | |
| 16S_F: 5’-GTGYCAGCMGCCGCGGTA-3’ | Earth Microbiome Project | 210570681 |
| 16S_R: 5’-GGACTACNVGGGTWTCTAAT-3’ | Earth Microbiome Project | 210570682 |
| Recombinant DNA | | |
| n/a |  |  |
| Software and Algorithms | | |
| Graphpad Prism 9 | Graphpad software | https://graphpad.com/scientific-software/prism |
| Empirical JTK_CYCLE | Hutchison *et al.* 2015 | https://github.com/alanlhutchison/empirical-JTK_CYCLE-with-asymmetry |
| QIIME2 | Hall and Beiko, 2018 | <https://qiime2.org> |
| ImageJ | Schindelin *et al.* 2015 | imagej.net |
| Adobe illustrator CC2020 | Adobe | https://www.adobe.com/product/photoshop.html |
| Orange3 | Demsar *et al.*, 2013 | <https://orangedatamining.com> |
| CircWave V1.4 | Roelof A. Hut *et al.* | <https://euclock.org/results/item/circ-wave.html> |
| Metascape | Zhou *et al.,* 2019 | <http://metascape.org> |
| Cytoscape v3.8.1 | Cline *et al.*, 2007 | https://cytoscape.org/ |
| STAR v2.6.1b | Dobin *et al.*, 2013 | https://github.com/alexdobin/STAR |
| DESeq2 | Love *et al.*, 2014 | https://github.com/mikelove/DESeq2 |
| fGSEA | Sergushichev *et al.,* 2016 | https://github.com/ctlab/fgsea |
| Other | | |
| JL Rat and Mouse/Auto 6F | LabDiet | Cat#5k67 |
| 0.1-mm-diameter zirconia/silica beads | BioSpec Products | Cat#11079101Z |
| Accu-Check Compact Plus Diabetes Monitoring Kit | Roche Diabetes Care | Cat#RO0510-01 |
| OneTouch Blood Glucose Monitoring System | OneTouch | Cat#024046 |
| Mini-PROTEAN TGX Stain-Free Gel | Bio-Rad | CAT#: 4568093 |
| Immobilon PVDF Transfer Membrane | Millipore | IPFL00010 |
| Heparin-coated microvette tubes | Sarstedt | Cat#16443100 |
| 6-well collagen-coated plates | Thermo Scientific | Cat#LS004188 |
| Accucheck Compack Test Strips | Roche Diabetes Care | Cat#FBA_TM82070 |
| OneTouch Ultra Blood Glucose Test Strips | OneTouch | Cat#LI020-994 |
